# Spatial patterning and immunosuppression of glioblastoma immune contexture in hypoxic niches

**DOI:** 10.1101/2022.03.01.482530

**Authors:** Anirudh Sattiraju, Sangjo Kang, Zhihong Chen, Valerie Marallano, Concetta Brusco, Aarthi Ramakrishnan, Li Shen, Dolores Hambardzumyan, Roland H Friedel, Hongyan Zou

## Abstract

Glioblastoma (GBM), a highly lethal brain cancer, is notorious for its immunosuppressive microenvironment, yet current immunotherapies are ineffective. Thus, understanding the immune contexture and governing factors of immunosuppression is crucial. Here, we identified a highly dynamic temporospatial patterning of tumor-associated macrophages (TAMs) corresponding to vascular changes in GBM: as tumor vessels transition from an initial dense regular network to later scant engorged vasculature, CD68^+^ TAMs shift away from perivascular regions to poorly vascularized areas. Remarkably, this process is heavily influenced by the immunocompetency of host animal, as tumor vessels in immunodeficient hosts remained dense and regular while TAMs evenly distributed. Utilizing a sensitive fluorescent reporter to track tumor hypoxia, we revealed that hypoxic niche controls immunosuppression by at least two mechanisms: first, attracting and sequestering activated TAMs in hypoxic zones, and second, reprograming entrapped TAMs towards an immunotolerant state. Indeed, entrapped TAMs also experience hypoxia and upregulate phagocytic marker *Cd68* and immunotolerant genes *Mrc1* and *Arg1*, thereby facilitating debris clearing, inflammatory containment, and immunosuppression in hypoxic zones. Mechanistically, we identified Ccl8 and IL-1β as two hypoxic niche factors released by TAMs in response to cues from hypoxic GBM cells, functioning to reinforce TAM retainment. Reciprocally, niche factors also shape the transcriptional responses of hypoxic tumor cells that exhibit quiescence and mesenchymal shift. Moreover, hypoxic niche factors are highly enriched in human GBMs, particularly mesenchymal subtype, and predict poor survival. Importantly, perturbing hypoxic niches resulted in reduced TAM sequestration and better tumor control. Together, understanding the mutual influence of immune contexture and metabolic landscape has important ramifications for improving efficacy of immunotherapies against GBM.

## INTRODUCTION

Glioblastoma (GBM), the most common primary brain cancer, remains highly lethal. A major contributing factor is its immunologically cold status, which poses significant hurdles for immunotherapy (Pombo Antunes et al., 2020). Although immune checkpoint inhibitors that stimulate cytotoxic T cells (e.g., anti-PD-1 and anti-CTLA-4) or chimeric antigen receptor (CAR) T cells have achieved some success in other solid cancers, they have not been effective for GBM patients (Brahm et al., 2020; Touat et al., 2020). Hence, new insights into GBM immune contexture and the governing factors of immunosuppression are urgently needed in order to improve the efficacy of immunotherapy.

Tumor-associated myeloid cells (TAMs) are a main component of the tumor microenvironment (TME) of GBM, and in extreme cases can constitute up to 50% of all cells in GBM (Klemm et al., 2020; Wei et al., 2020). It has been shown that TAMs can be re-educated by niche cues to support GBM growth, promote angiogenesis, and dampen T cell activation (Andersen et al., 2021). Current strategies to target TAMs by either reducing their numbers or reversing their pro-tumorigenic state have provided promising results in animal GBM models (Akkari et al., 2020; Pyonteck et al., 2013), but have not yet yielded significant clinical gain for GBM patient survival (Bejarano et al., 2021; Butowski et al., 2016). Thus, mapping the spatial organization of TAMs and dissecting contextual cues of the TME will facilitate the development of new strategies to ablate or reprogram TAMs.

A drawback of the commonly used methods to study the immune cell composition during GBM progression, e.g., flow cytometry and single cell RNA-sequencing (scRNA-seq), is the lack of spatial information. As niche cues from the TME play instructive roles on gene expression and functional specification, we reasoned that mapping the geographic patterning of immune cells may provide new insights into the mechanisms and driving forces of immunosuppression in GBM.

To delineate the spatial relationship of TAM patterning and metabolic state of the TME during GBM progression, developing a faithful reporter for activation of hypoxia-induced transcription factor (HIF), a master regulator of physiological adaptation to low oxygen (Weidemann and Johnson, 2008) is crucial. Current approaches to detect tumor hypoxia using compounds such as pimonidazole and EF5 (Koch et al., 1995; Varia et al., 1998) or PET imaging (Stokes et al., 2016; Verhoeven et al., 2019) are limited by low cellular resolution, incomplete tissue penetration, and lack of a direct link to HIF activity. Conventional genetic reporters with HIF-responsive elements (HRE) driving the expression of green fluorescent protein (GFP) are hampered by the requirement of O_2_ for fluorophore maturation. We thus took advantage of a novel fluorescent protein UnaG, which does not require O_2_ to become fluorescent (Kumagai et al., 2013). Indeed, HRE-UnaG reporter can faithfully track HIF activity in hypoxic cells (Erapaneedi et al., 2016). We engineered a lentiviral HRE-UnaG vector for stable transduction of GBM cells for in vivo tracking of HIF activity during GBM progression.

We charted temporospatial organization of TAMs in a murine orthotopic GBM model in a fully immunocompetent background. This highly dynamic process involves active cell-cell sorting and development of pseudopalisading patterns. Mechanistically, we identified Ccl8 and IL-1β as two hypoxic niche factors released by TAMs in response to instructive cues from hypoxic GBM cells, functioning to promote TAM trafficking and sequestration in hypoxia zones, while IL-1β is also identified as an upstream regulator of the transcriptional responses of hypoxic GBM cells. Additionally, the HRE-UnaG HIF reporter allowed us to capture the in vivo GBM hypoxia gene signature, which displayed quiescence and mesenchymal features, as well as a high degree of immune signaling, in line with an active role of hypoxic GBM cells in attracting and reprogramming TAMs. The hypoxic niche factors are highly represented in human recurrent GBMs and predict poor survival. Finally, we demonstrated that strategies to perturb hypoxic niches reduced TAM sequestration and improved tumor control.

## RESULTS

### Dynamic temporospatial patterning of TAMs and a critical role of immune status of GBM host

To characterize the spatial organization of TAMs in a fully immunocompetent microenvironment, we utilized the orthotopic murine GL261 GBM model in syngeneic C57BL/6 wildtype (B6-WT) mouse host. At 4 weeks post-transplant, immunofluorescence (IF) for the myeloid marker Iba1 revealed a highly heterogenous pattern of TAMs (**Fig. 1a**). Similar spatial patterns were also observed for TAMs expressing macrophage antigen F4/80 or integrin α4 (CD49d), a marker of blood-borne monocyte-derived macrophages (MDMs) (Bowman et al., 2016) (**Fig. 1a; Fig. S1a**). Notably, microglia-derived TAMs labeled by Tmem119 resided mostly at GBM periphery, while MDM-derived TAM (integrin α4^+^) congregated mostly in GBM interior (**Fig. S1b**). Further IF staining and quantifications confirmed the distinct spatial organization of TAMs expressing either phagocytic marker CD68 or pro-tumor/immunosuppressive/anti-inflammatory (M2) marker mannose receptor Mrc1 (CD206) (**Fig. 1b**).

**Figure 1.**
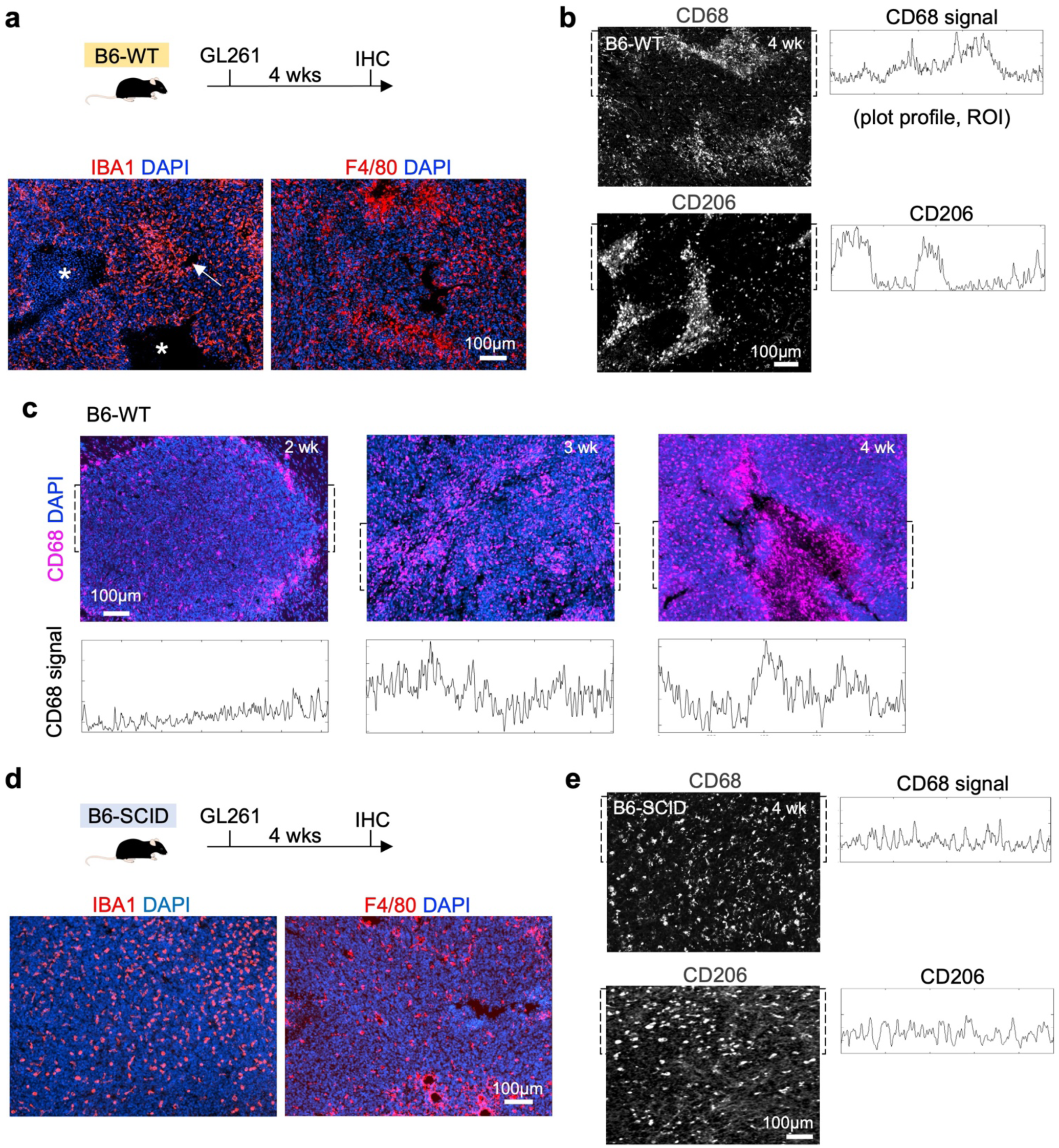
Dynamic temporospatial patterning of TAMs and the influence of immune status of GBM. **(a)** GL261 GBM cells were orthotopically transplanted in C57BL/6 wild-type (B6-WT) host mice. IF images show distinct spatial patterning of IBA1^+^ and F4/80^+^ TAMs in GBM interior. **(b)** IF images and quantification plots of relative signal intensity of CD68 or CD206 within the regions of interest (ROI, dashed brackets) show distinct spatial patterning of TAMs in GL261 GBM. **(c)** IF images and relative signal intensities of CD68 within ROIs (dashed brackets) at different time points show increased spatial patterning of CD68^+^ TAMs as GL261 progresses. **(d)** GL261 tumors were orthotopically established in C57BL/6 immunodeficient (B6-SCID) host mice. IF images show relatively uniform distribution of TAMs (IBA1^+^ or F4/80^+^) in GBM interior. **(e)** IF images and quantification plots of relative signal intensity within ROIs show even distribution of CD68^+^ and CD206^+^ TAMs in GL261 GBM in B6-SCID host.

Interestingly, the spatial patterning of TAMs appeared highly dynamic during GBM expansion: at early stage (2 weeks post-transplant), CD68^+^ TAMs largely accumulated at tumor periphery, while in tumor interior they were relatively evenly distributed; but by 3-4 weeks post-transplant, they became increasingly congregated in distinct zones inside GBM (**Fig. 1c**).

As immunodeficient mice are commonly utilized for patient-derived xenotransplant (PDX) models, we wondered whether the immune status of the host might affect the spatial arrangement of TAMs. We thus repeated the study of intracranial transplantation of GL261 in C57BL/6-SCID mice (B6-SCID) lacking functional T and B lymphocytes. Interestingly, we observed a more uniform distribution of TAMs at all stages of GBM progression, including TAMs positive for Iba1, F4/80, CD68, or CD206 (**Fig. 1d, e; Fig. S1c**). We also confirmed these findings with the commonly used ICR-SCID outbred immunodeficient mice, which again showed a relatively uniform distribution of TAMs labeled by Iba1 and CD68 (**Fig. S1d**). Together, these results revealed a highly dynamic geographic patterning of TAMs during GBM progression, which is strongly influenced by host immune status.

### TAM patterning parallels vascular changes during GBM progression

As TAMs in the GBM interior are mainly composed of blood-borne MDMs, we wondered if TAM patterning might be related to vascular rearrangements during GBM progression. To this end, we first compared the vascular architecture of GL261 tumors established in B6-WT vs. B6-SCID hosts and found a pronounced difference (**Fig. 2a**). In B6-WT hosts, the tumor vessels (PECAM1^+^) showed a progressive transformation from an initial dense regular network at 2 weeks post-transplant to a later sparse, engorged, and tortuous vasculature at 4 weeks post-transplant (**Fig. 2a-c**). In contrast, in B6-SCID hosts, tumor vascular networks remained dense with regular branching patterns and uniform lumen size throughout GBM progression (**Fig. 2a, b, d**). This indicates that a fully functional adaptive immunity in B6-WT hosts plays a critical role in the aberrant vascular development during GBM expansion.

**Figure 2.**
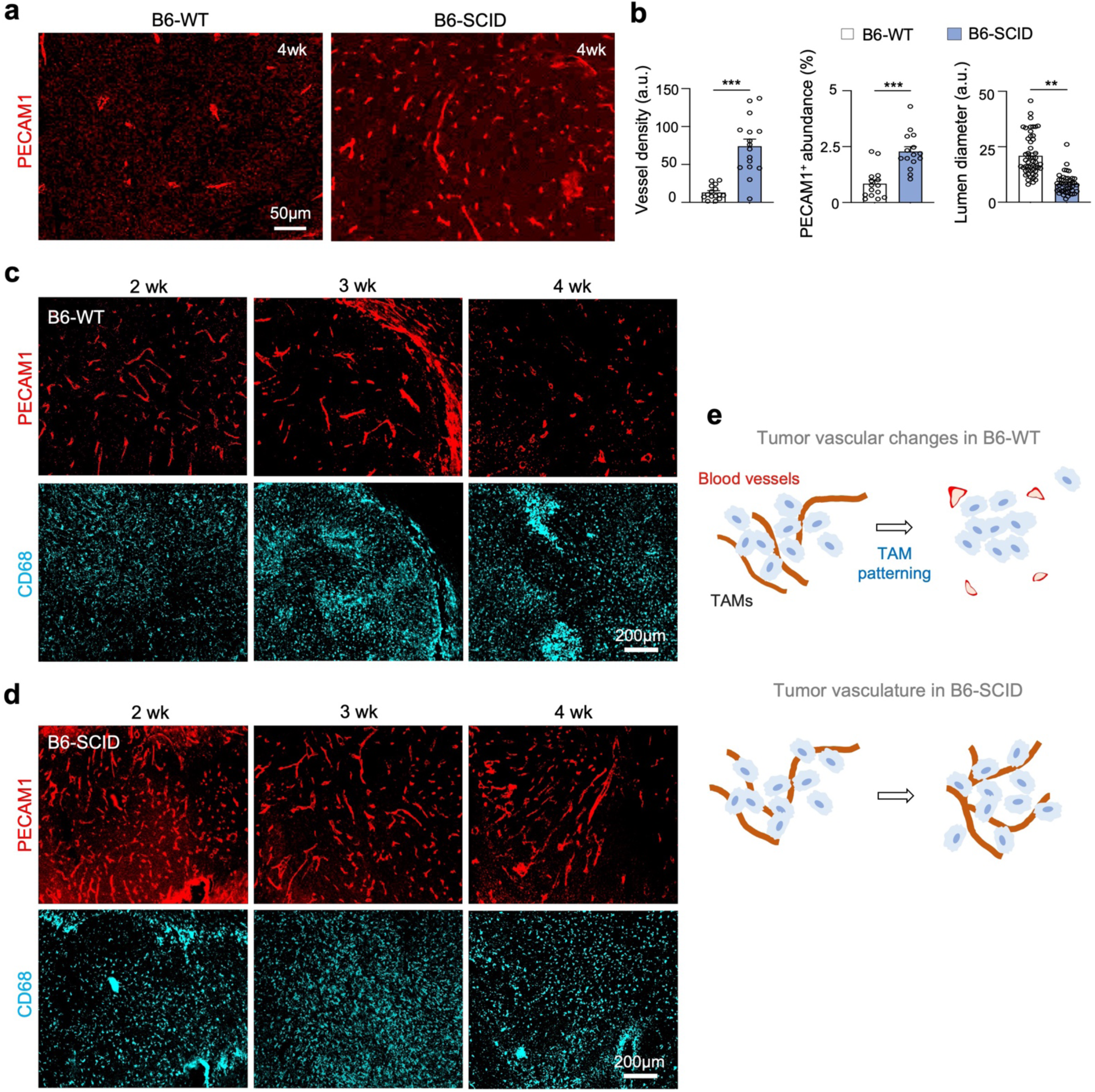
TAM patterning parallels vascular changes during GBM progression. **(a, b)** IF images and quantifications reveal more robust tumor vasculature (PECAM1^+^) and less vessel engorgement in tumors established in B6-SCID mice as compared to B6-WT host at 4 weeks post-transplant. Quantifications of random tumor areas from 5 mice per group, unpaired t-test, \*\**P*<0.01, \*\*\**P*<0.001. **(c)** IF images of GL261 GBM in B6-WT hosts show temporospatial confinement of CD68^+^ TAMs in distinct zones that parallels progressive vascular changes from dense regular network to sparse, tortuous, and engorged vessels. **(d)** IF images of GL261 GBM in B6-SCID hosts show consistent tumor vasculature throughout tumor progression, and in parallel, CD68^+^ TAMs are uniformly distributed without geographic patterning. **(e)** Diagrams depicting TAM patterning corresponding to vascular changes that are influenced by host immune status.

Correspondingly, in GBM developed in B6-WT hosts, the striking vascular alterations were paralleled by progressive changes of TAM patterning: at early stages (2-3 weeks post-transplant), CD68^+^ cells mainly clustered around blood vessels, consistent with transvascular influx of MDMs into GBM; by 3-4 weeks post-transplant, with increasingly sparse blood vessels, CD68^+^ cells became congregated in poorly vascularized areas (**Fig. 2c**). By contrast, in B6-SCID hosts, while blood vessels remained dense and regular throughout GBM progression, CD68^+^ TAMs remained evenly distributed (**Fig. 2d**). Hence, the spatial organization of CD68^+^ TAMs appeared closely associated with the vascular alterations during GBM expansion, and this process is heavily influenced by the immune status of host animals (**Fig. 2e**).

### Tracking the emergence of tumor hypoxic zones with HRE-UnaG reporter

As inadequate blood supply of tumor tissues would lead to hypoxia, we next asked if the spatial patterning of active CD68^+^ TAMs may be caused by the emergence of hypoxic zones during GBM progression. To investigate this hypothesis, we introduced the HIF reporter HRE-UnaG by stable lentiviral transduction into GL261 GBM cells (**Fig. 3a**). The HRE-UnaG reporter has been shown to faithfully report HIF activity in hypoxic cells (Erapaneedi et al., 2016), and we validated first in vitro the robust induction of UnaG in GL261 cells upon exposure to hypoxia (1% O_2_) and a rapid turnover upon return to normoxia (UnaG is fused with a PEST degron for fast temporal dynamics) (**Fig. S2a**).

**Figure 3.**
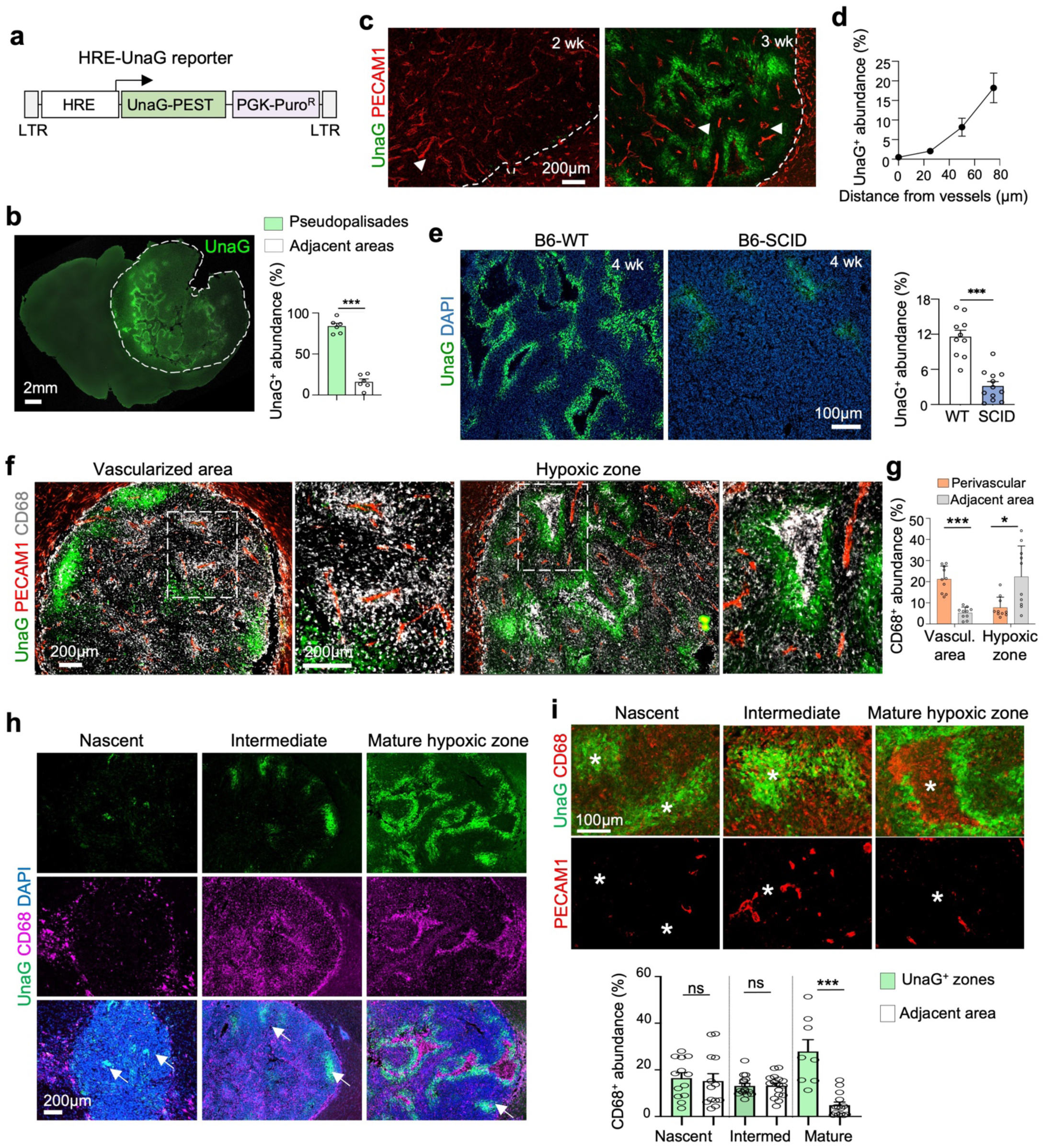
Emergence of hypoxic zones, TAM trafficking, and development of pseudopalisades. **(a)** Diagram of lentiviral HRE-UnaG reporter carrying a HIF sensitive promoter driving the expression of UnaG fused to a PEST degron for fast turnover. See Methods for details. **(b)** GL261-HRE-UnaG GBM established in B6-WT host harbored abundant UnaG^+^ tumor cells. Quantification shows that majority of UnaG^+^ cells resided in pseudopalisading area. n=4 mice per group, unpaired t-test; \*\*\**P*<0.001. **(c)** IF images show the emergence of UnaG^+^ tumor cells between 2-3 weeks post-transplant in poorly vascularized areas. Note reduced vascular density and engorged lumen (arrowheads) during tumor progression. **(d)** Quantification confirms increased UnaG abundance with increasing distance from blood vessels. n=11 fields for each distance. **(e)** IF images and quantification reveal much smaller hypoxic zones in GL261 GBM established in B6-SCID hosts. n= 10 areas, unpaired t-test. ***, *P*<0.001. **(f, g)** IF images and quantification demonstrate a transition of CD68^+^ TAMs from perivascular region to hypoxic zones in GL261 GBM, coinciding with vascular transformation. Enlarged images of boxed areas are shown on the right. Note that entrapped CD68^+^ TAMs were corralled by UnaG^+^ tumor cells in pseudopalisading patterns. n = 10 areas for each group; multiple unpaired t-tests; \**P*<0.05; \*\*\**P*<0.001; ns, not significant). **(h)** IF images show development of hypoxic zones (UnaG^+^) from nascent (arrows) to mature psuedopalisades, corresponding to vascular changes (PECAM1^+^) during GL261 progression. **(i)** IF images show a gradual cell-cell sorting of CD68^+^ TAMs and UnaG^+^ tumor cells in hypoxia areas (asterisks) during the transition from nascent hypoxic zones to mature pseudopalisades, coinciding with aberrant vascular development (PECAM1^+^). Quantification below shows enrichment of CD68^+^ cells in mature hypoxic zones. n=3 mice per group, One-way ANOVA, \*\*\**P*<0.001; ns, not significant.

We then transplanted GL261-HRE-UnaG cells intracranially into B6-WT mice, and at 4 weeks post-transplant, we found abundant UnaG^+^ (HIF^ON^) cells inside the tumor in pseudopalisading patterns (**Fig. 3b**). Time course analyses indicated that UnaG^+^ cells emerged between 2-3 weeks post-transplant in areas distant from blood vessels (**Fig. 3c, d**). The UnaG^+^ cells indeed expressed high levels of glucose transporter 1 (GLUT1), a direct HIF target gene (Harris, 2002), confirming faithful reporting for HIF activity in vivo (**Fig. S2b**). The spatial arrangement of hypoxic zones (UnaG^+^ or GLUT1^+^) in GL261 tumors closely resembled the GLUT1^+^ patterns in human GBM (**Fig. S2b**). Quantifications demonstrated that the abundance of UnaG^+^ areas and the size of pseudopalisading zones increased with tumor progression, while the density of tumor vasculature decreased but vessel lumen engorged (**Fig. S2c-e**).

We next compared HRE-UnaG reporter with Pimonidazole (Pimo), a frequently used hypoxia-sensitive compound that is covalently linked to proteins by cellular reductases under low oxygen (Varia *et al*., 1998). We found that HRE-UnaG labeled more tumor cells in wider areas than Pimo, especially in nascent hypoxic zones populated with small UnaG^+^ but Pimo^-^ aggregates, whereas in more mature hypoxic zones with pseudopalisading patterns, UnaG largely overlapped with Pimo (**Fig. S2f**). These data support a higher sensitivity of HRE-UnaG in unveiling nascent hypoxic zones, in line with its capability to report HIF activity below 5% O_2_ tension (Erapaneedi *et al*., 2016; Schmitz et al., 2020), while Pimo mostly labels hypoxic cells below 1% O_2_ tensions (Young and Möller, 2010).

Given the marked differences of tumor vasculature in GBM growing in B6-WT vs. B6-SCID hosts, we next compared the UnaG expression patterns in these two GBM paradigms. We indeed observed a striking difference: whereas UnaG^+^ GBM cells were highly abundant in B6-WT hosts by 3-4 weeks post-transplant, they remained scant in B6-SCID hosts even though tumors had reached larger sizes at matching time points (**Fig. 3e; Fig. S3a, b**). Interestingly, temporal analyses revealed numerous individual UnaG^+^ cells scattered throughout tumor tissues at 2-3 weeks post-transplant in B6-SCID hosts, likely reflecting fast tumor expansion outpacing vascular development, but by 4 weeks post-transplant, they were largely resolved (**Fig. S4**), indicating robust tumor angiogenesis and reoxygenation in immunodeficient hosts. Hence, sustained increase of tumor hypoxia is not just dictated by rapid tumor expansion outstripping vascular supply, but also influenced by the immune status of host animals affecting vascular development.

### The spatial arrangement of TAMs is linked to pseudopalisade development and hypoxic zone maturation

We next applied the HRE-UnaG HIF reporter to delineate the spatial relationship of the immune landscape with tumor hypoxia during GBM progression. We found that at early stages between 2-3 weeks post-transplant into B6-WT hosts, UnaG^+^ tumor cells were less abundant and only present in nascent pseudopalisading areas, while CD68^+^ TAMs mostly accumulated around blood vessels (**Fig. 3f-h**). However, at later stages between 3-4 weeks post-transplant, CD68^+^ TAMs were attracted to and later confined in hypoxic zones surrounded by a rim of UnaG^+^ tumor cells (**Fig. 3f-h**). Hence, the shift of CD68^+^ TAM localization from perivascular regions towards hypoxic cores coincided with the expansion of hypoxic zones and maturation of pseudopalisades.

The sensitivity and high cellular resolution conferred by the HRE-UnaG reporter allowed us to appreciate in detail the dynamic development of hypoxic zones and pseudopalisades, involving the recruitment of CD68^+^ TAMs, cell-cell sorting, and corralling of TAMs by UnaG^+^ GBM cells **(Fig. 3h, i)**. These changes may facilitate debris clearing of necrotic cells and inflammatory containment, thus contributing to an immunosuppressive status of the microenvironment.

### Confined TAMs experience hypoxia and express immunotolerance markers

We suspected that the entrapped TAMs may also experience hypoxia. Indeed, co-staining showed an overlap of CD68 and CD206 with hypoxia markers Glut1 and Pimo, respectively (**Fig. 4a, b**). To further examine the functional state of these hypoxic TAMs, we conducted additional IF, which showed that the entrapped TAMs not only expressed CD206, but also Arginase 1 (Arg1), both being immunotolerance markers (**Fig. 4c, d**).

**Figure 4.**
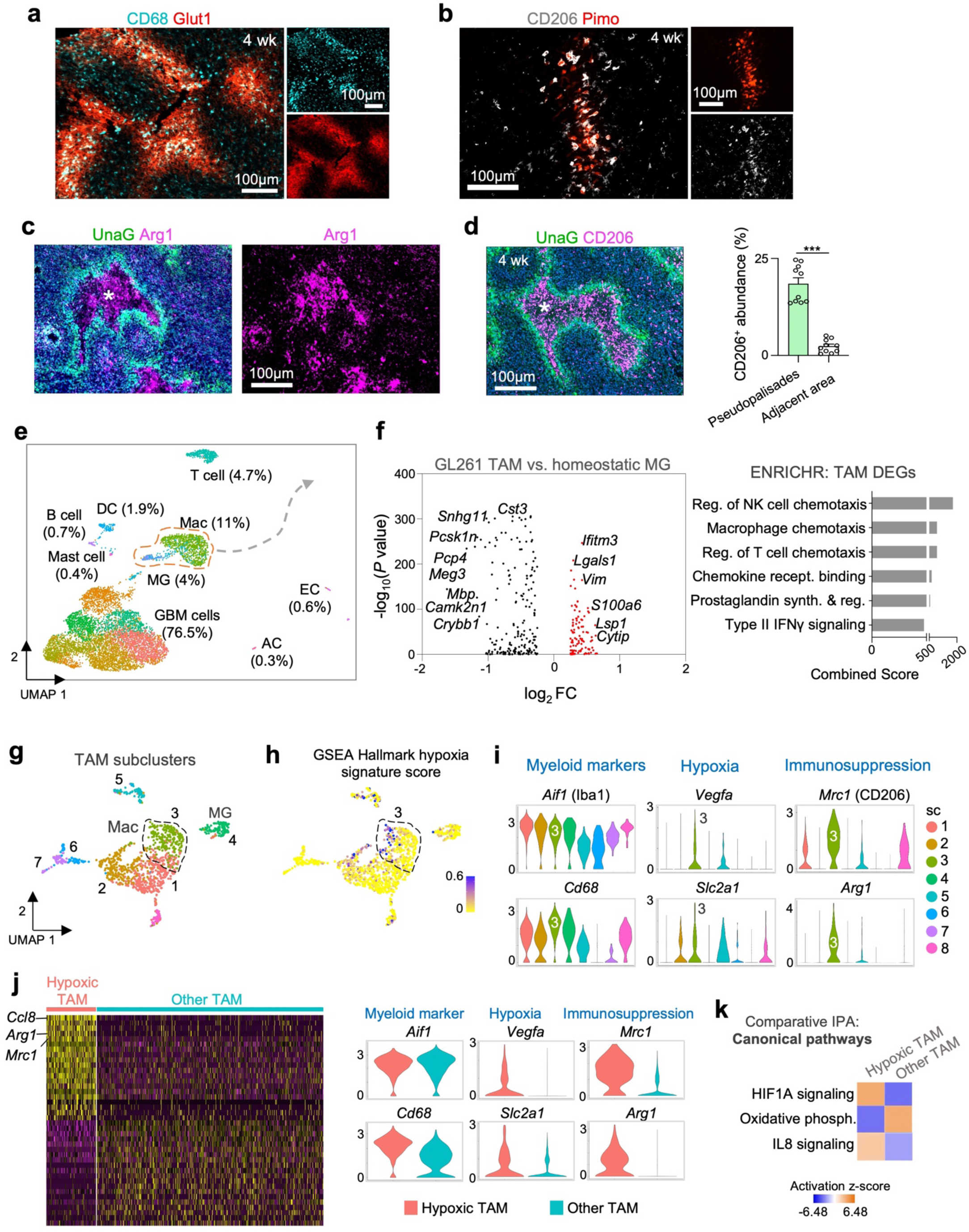
Entrapped TAMs experience hypoxia and express immunosuppressive markers. **(a, b)** IF images show overlap of CD68 and CD206 with hypoxia marker GLUT1 or Pimo, respectively. **(c, d)** IF images show that sequestered TAMs (asterisks) highly express immunotolerance markers Arginase-1 (Arg1) and CD206. Quantification of n=10 randomly selected area from 3 mice per condition, unpaired t-test, \*\*\**P*<0.001. **(e)** UMAP plot of scRNA-seq data from GL261 GBM in B6-WT host at 4 weeks revealed GBM cells (occupying the largest share) and 8 major stromal cell types. **(f)** Left, volcano plot shows differentially expressed genes (DEGs) in GL261 TAMs (microglia and macrophages combined) as compared to homeostatic microglia (data from Zeisel et al., 2018). Right, ENRICHR analysis shows that the DEGs in TAMs are enriched for genes involved in macrophage chemotaxis and regulation of T cell chemotaxis, in addition to chemokine receptor signaling and immune modulation. **(g)** UMAP plot of TAMs shows eight subclusters: one for microglial (MG) and seven for macrophages (Mac). **(h)** UMAP plot of TAMs shows enrichment scores for Hallmark hypoxia gene signature, with TAM subcluster 3 displaying the strongest enrichment. **(i)** Violin plots show relative expression of the indicated marker genes in TAM subclusters. Note that subcluster 3 TAMs highly express *Cd68*, classic hypoxia-response genes (*Slc2a1*, *Vegfa)* and immunosuppression markers (*Mrc1*, *Arg1)*. **(j)** Heatmap reveals DEGs in hypoxic TAMs (subcluster 3) relative to all other TAMs based on scRNA-seq of GL261 GBM, with *Ccl8* being the top upregulated DEG. Violin plots confirm high expression of the indicated markers in hypoxic TAM. **(k)** Ingenuity Pathway Analysis (IPA) shows top canonical pathways enriched in hypoxic TAM (subcluster 3) compared to other TAM subclusters.

To better define transcriptional profiles of TAMs, in particular the hypoxic TAM population, we performed scRNA-seq on GL261-HRE-UnaG GBM at 4 weeks post-transplant using the 10X Genomics platform (**Fig. 4e**). Cluster analysis revealed 9 major cell types, with GBM cells occupying the largest share (76.5%), followed by tumor-associated macrophages (11%) and microglia (4%), other immune cells (dendritic cells 1.9%, T cells 4.7%, B cells 0.68%, mast cells 0.38%), endothelial cells (0.6%), and a small population of astrocytes (0.3%) (**Fig. S5**).

We first compared TAMs in GL261 GBM with homeostatic microglia in normal brain (data set from (Zeisel et al., 2018)). We found that the differentially expressed genes (DEGs) in TAMs are highly enriched for pathways linked to macrophage chemotaxis, regulation of chemotaxis of natural killer cells or T cells, oxidative damage, prostaglandin synthesis, and type II interferon (IFN) signaling **(Fig. 4f)**. Hence, TAMs in GBM launch gene programs for chemotaxis, immunomodulation, and metabolic adaptation.

The TAM population in GL261 GBM can be further partitioned into eight subclusters (sc), with microglia (sc-4) clearly distinguishable from macrophages (**Fig. 4g**). Among the seven macrophage subclusters, sc-3 TAMs are enriched for GSEA hallmark hypoxia genes (**Fig. 4h**), and they express higher levels of hypoxia marker genes *Vegfa* and *Slc2a1* (encoding GLUT1), immunotolerance genes *Mrc1* (CD206) and *Arg1,* as well as phagocytic marker *Cd68,* while *Iba1* is more ubiquitously expressed among TAM subclusters (**Fig. 4i; Fig. S6a, b**). Sc-3 TAMs therefore likely correspond to the hypoxic TAMs entrapped in UnaG^+^ zones observed by histological analyses. A pseudotime analysis suggested that sc-3 TAMs are differentiated from *Ccr2*^+^ monocytes (sc-2) (**Fig. S6c**).

We further compared the transcriptomes of sc-3 TAMs relative to other TAMs (**Fig. 4j**). Results confirmed that *Cd68, Slc2a1*, *Mrc1,* and *Arg1* are among the top DEGs in hypoxic TAMs. Ingenuity Pathway Analysis (IPA) further demonstrated HIF1A signaling and IL8 signaling as the top canonical pathways enriched in sc-3 TAMs (**Fig. 4k**). These results provided transcriptomic evidence for a specific hypoxic and immunosuppressive TAM population in GBM.

### Sequestration and immune exhaustion of cytotoxic T cells in hypoxic zones of GBM

Aside from the spatial confinement of TAMs in pseudopalisading areas, CD8a^+^ cytotoxic T lymphocytes (CTLs) were also found to be sequestered in UnaG^+^ zones; whereas FoxP3^+^ regulatory T cells (Tregs) were abundant in UnaG^-^ tumor areas (**Fig. S6d-e**). We further analyzed transcriptomes of the T cells in our scRNA-seq data, which revealed 6 subclusters among *Cd3e+* (CD3) lymphocytes, with sc-1, 2, and 6 annotated as CTLs, sc-3 as Tregs, sc-4 as natural killer (NK) cells, and sc-5 as CD4 T helper cells, based on the expression of cell type-specific markers (**Fig. S6f**). Among the CTLs, the largest subcluster (CTL-a) was associated with GSEA for glycolysis, hypoxia, as well as immune checkpoint blockade transcriptional programs (**Fig. S6g-l**). CTL-a also showed high expression of T cell exhaustion marker *Havcr2* (**Fig. S6k**), consistent with earlier finding that the CTLs arriving in GBM become mostly exhausted (Broekman et al., 2018; Watson et al., 2021). Together, the histological and transcriptomic data indicate entrapment of CTLs in pseudopalisading zones, with signs of hypoxia and immune exhaustion, thus further linking tumor hypoxia to immunosuppression.

### Progressive spatial confinement of TAMs involves Ccl8/12 and IL-1β

We next explored the molecular mechanisms underlying TAMs trafficking and sequestration in pseudopalisading areas. We first focused on cytokine *Ccl8*, which was one of the top upregulated genes in hypoxic (sc-3) TAMs (see Fig. 4j; **Fig. 5a; Fig. S7a**). We hypothesized that hypoxic GBM cells may send instructive cues to induce the expression of Ccl8 in TAMs, which in turn would promote their trafficking to hypoxic zones and development of pseudopalisades. First, scRNA-seq data showed that the Ccl8 receptors *Ccr1* and *Ccr5* are highly expressed in TAMs, indicating responsiveness of TAMs to Ccl8 (**Fig. S7a**). Second, to test whether exposure to hypoxic tumor cells would induce *Ccl8* in TAMs, we exposed primary bone marrow-derived macrophages (BMDMs) to conditioned media (CM) from GL261 cells cultured either in hypoxic or normoxic conditions. Strikingly, CM from GL261 cells resulted in a marked upregulation of *Ccl8* in BMDMs, with the CM from hypoxic GL261 cells exerting a stronger effect (**Fig. 5b**). Notably, the immune genes *Cd68* and *Arg1* were also robustly induced in BMDMs by CM of hypoxic GL261 GBM cells (**Fig. S7b**).

**Figure 5.**
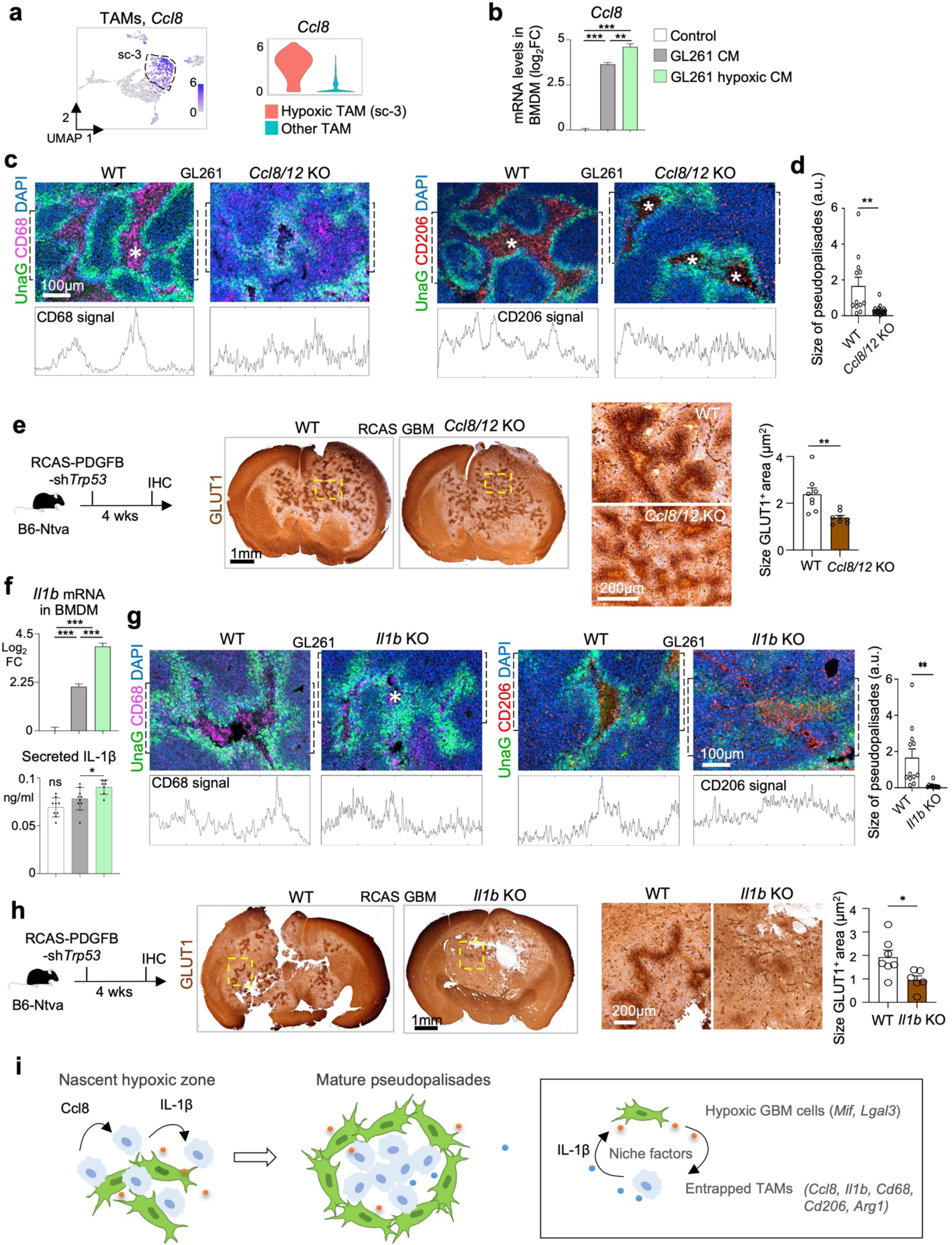
Progressive spatial confinement of TAMs in hypoxic zones requires Ccl8/12 and IL-1β. **(a)** Left, UMAP plot shows high expression of *Ccl8* in TAM subcluster 3 (sc-3). Right, violin plot confirms higher expression of *Ccl8* in hypoxic TAMs compared to other TAMs. **(b)** qRT-PCR results show upregulation of *Ccl8* in BMDMs exposed to conditioned media (CM) from GL261 cells for 24 hrs. The CM was prepared from GL261 cells cultured in 1% O_2_ or normoxia conditions. n=3 wells per condition, one-way ANOVA, \*\**P*< 0.01, \*\*\**P*<0.001, ns, not significant. **(c, d)** IF images and quantifications of signal intensity of CD68 and CD206 in ROI (dashed box) reveal that GL261 GBM transplanted in *Ccl8/12* KO host mice harbored smaller pseudopalisades and displayed less well-defined entrapment of TAMs. n=3 mice per group, unpaired t-test; \*\**P*<0.01. **(e)** RCAS GBM (diagram on left) generated in *Ccl8/12* KO mice harbored smaller GLUT1^+^ pseudopalisades than RCAS GBM in WT controls. n=8 hypoxic zones per genotype, unpaired t-test; \*\**P*< 0.01. **(f)** qRT-PCR and ELISA revealed higher levels of *Il1b* mRNA and secreted IL-1ß protein from cultured bone marrow-derived macrophages treated with CM from GL261 cells (cultured in 1% O_2_ or normoxia) relative to control media. n=3 wells per condition, one-way ANOVA; \**P*< 0.05, \*\*\**P*<0.001, ns, not significant. **(g)** IF image and quantifications of signal intensity of CD68 and CD206 in ROI (dashed box) reveal that that GL261 GBM transplanted into *Il1b* KO host mice harbored smaller pseudopalisades and less well-defined entrapment of TAMs than in WT hosts. n=3 mice per group, unpaired t-test; \*\**P*<0.01. **(h)** Representative immunohistochemistry images show that RCAS GBM generated in *Il1b* KO mice harbored smaller GLUT1^+^ pseudopalisades than RCAS GBM in WT controls. n=7 hypoxic zones in WT and n=6 hypoxic zones in *Il1b* KO animals, unpaired t-test; \**P*<0.05). (**i**) Diagram of hypoxia-induced immunosuppression by two mechanisms: i) hypoxic tumor cells launch a program to attract and sequester TAMs through induction of Ccl8 and IL-1ß, and ii) entrapped TAMs experience hypoxia and are reprogrammed to express CD68 and immunosuppressive markers CD206 and Arg1.

To assess the functional significance of *Ccl8* upregulation for TAM patterning, we established GL261 GBM in *Ccl8/12* knockout (KO) mice (Tsou et al., 2007). Of note, this KO mutation affects *Ccl8* and *Ccl12* due to their close genomic proximity; however, *Ccl12* is not induced in hypoxic TAMs (**Fig. S7c**), and it is absent from human genome, indicating a lesser role for hypoxia-regulated TAM patterning. We found that while UnaG^+^ tumor cells were abundantly detected in both *Ccl8/12* KO and control hosts, the UnaG^+^ pseudopalisades were significantly smaller in *Ccl8/12* KO hosts. Correspondingly, the spatial patterns of CD68^+^ or CD206^+^ TAMs also appeared less well-developed in KO mice (**Fig. 5c, d**).

To further corroborate the role of Ccl8 in governing the spatial organization of TAMs in hypoxic zones and the development of pseudopalisades, we also studied the effect of *Ccl8/12* KO with the autochthonous mouse RCAS GBM model, which is based on GBM induction by an avian viral system expressing GBM-driving oncogenes or immunosuppressors (Herting et al., 2019). At 30 days after induction of RCAS-PDGFB-sh*Trp53* GBM in either wild-type controls or *Ccl8/12* KO mice, we examined the development of hypoxic zones (GLUT1^+^) (**Fig. 5e**). Similar to the findings in GL261 tumors, the RCAS GBMs established in *Ccl8/12* KO mice contained significantly smaller hypoxic zones and less developed pseudopalisades than controls (**Fig. 5e**). Hence, both GL261 and RCAS GBM models demonstrated that *Ccl8/12* deficiency in stromal cells compromised TAM patterning and maturation of hypoxic pseudopalisades.

Interestingly, we also found that interleukin-1 beta (*Il1b*) mRNA was highly induced in BMDMs when exposed to CM from GL261 cells, and more so by CM from hypoxic GL261 cells (**Fig. 5f**). As activated macrophages secrete IL-1β after the proteolytic cleavage of pro-IL-1β (Herting *et al*., 2019), we performed ELISA on BMDM culture supernatants, which confirmed that CM from hypoxic GL261 cells significantly increased IL-1β secretion (**Fig. 5f**). As IL-1β is a major cytokine that modulates inflammatory responses and cytotoxicity (Gabay et al., 2010), we investigated its contribution to the development of hypoxic zones and spatial patterning of TAMs. First, scRNA-seq data showed that *Il1b* was specifically expressed in TAMs in GL261 (**Fig. S7d**). We then established GL261-HRE-UnaG tumors in *Il1b* KO mice and wild-type controls, which revealed that hypoxic zones outlined by UnaG^+^ tumor cells were much smaller in *Il1b* KO hosts, with reduced entrapment of CD68^+^ and CD206^+^ TAMs in hypoxic zones (**Fig. 5g**).

We next corroborated the finding that IL-1β controls the TAMs trafficking and maturation of hypoxic pseudopalisades by analyzing RCAS GBM in *Il1b* KO vs. wild-type mice (Herting *et al*., 2019). We observed significantly smaller hypoxic zones (GLUT1^+^) and near absent pseudopalisading patterns in RCAS GBM developed in *Il1b* KO as compared to wild-type hosts (**Fig. 5h**). Together, these results establish Ccl8 and IL-1β as TAM-derived niche factors that critically control immune landscape and pseudopalisade development. It is noteworthy that in the RCAS GBM studies with KO host mice, *Ccl8* or *Il1b* were absent from both tumor and non-tumor cells, whereas in GL261 GBM transplant models, genes were deleted only in non-tumor cells, which may account for the more pronounced spatial perturbances of hypoxic zones in the RCAS GBM models.

In aggregate, our data support the working model that hypoxic GBM cells release factors to induce upregulation of cytokines Ccl8 and IL-1β in TAMs, which in turn promote trafficking and sequestration of TAMs in hypoxic zones, facilitating TAM entrapment and pseudopalisade development (**Fig. 5i**).

### In vivo gene signature of hypoxic GBM cells reveals a high degree of immune signaling

The HRE-UnaG reporter provided a unique opportunity to capture the physiological in vivo gene signature of HIF^ON^ GBM cells, as these cells can be easily distinguished from HIF^OFF^ counterparts by the *UnaG* transcript levels. Examination of our scRNA-seq data showed that the UnaG^+^ (HIF^ON^) population formed a separate subcluster (sc-5) from UnaG^-^ cells, signifying a distinct transcriptional state (**Fig. 6a; Fig. S8a, b**). Indeed, UnaG^+^ cells exhibited a mesenchymal shift, as evidenced by a higher MES2 signature score (Neftel et al., 2019) (**Fig. 6a**). This echoes a recent large-scale scRNA-seq study on human GBMs showing that among the different states of GBM cells, the MES-like state is most associated with a hypoxia signature (Neftel *et al*., 2019).

**Figure 6.**
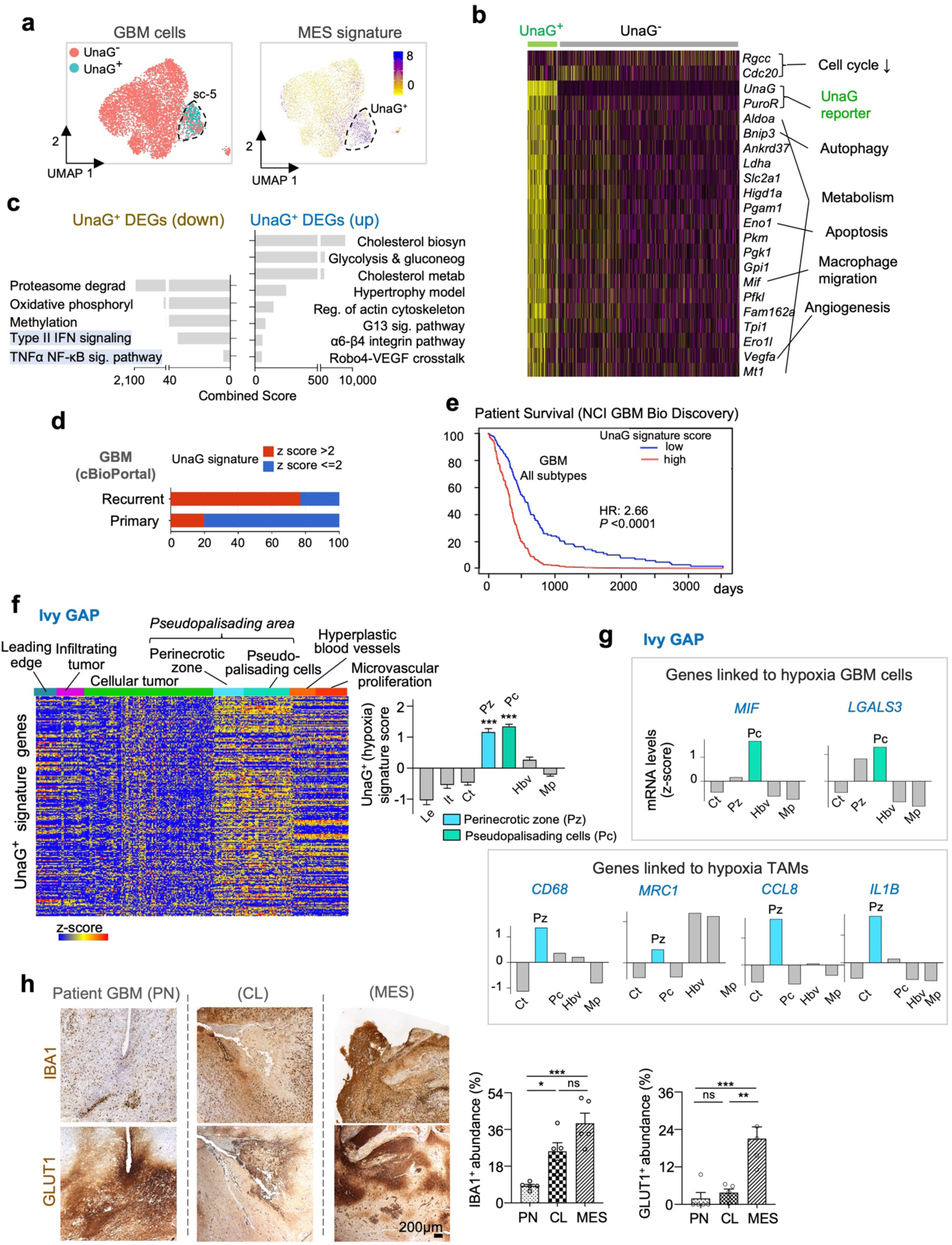
In vivo UnaG gene signature predicts poor outcome and is enriched in pseudopalisading areas in human GBM. **(a)** UMAP plot shows that UnaG^+^ cells occupy a subcluster (SC-5) distinct from UnaG^-^ population based on scRNA-seq data from GL261 GBM. UnaG^+^ subcluster was enriched for mesenchymal (MES) gene signature. **(b)** Heatmap shows top DEGs in UnaG^+^ GBM cells compared to UnaG^-^ counterparts. Annotations of main functions of DEGs are shown on the right. **(c)** Pathway enrichment analyses of upregulated and downregulated DEGs of UnaG^+^ GBM cells by ENRICHR (using WikiPathways 2019 gene sets). **(d)** The UnaG^+^ GBM gene signature is enriched in recurrent human GBM (cBioPortal, TCGA GBM data). **(e)** The UnaG^+^ GBM gene signature correlates with poor patient survival (NCI BioDiscovery portal, TCGA GBM data). **(f)** Heatmap shows higher expression of UnaG^+^ gene signature genes in anatomical domains of peri-necrotic zone (PZ) and pseudopalisading cells (PC) as compared to other areas based on analysis of human GBM specimens (Ivy GBM Atlas Project, Ivy GAP). The graph to the right shows summarized values of the heatmap. One-way ANOVA, \*\*\**P*<0.001. **(g)** Graphs based on Ivy GAP data show that hypoxia niche genes associated either with UnaG^+^ tumor cells (top) or entrapped TAMs (bottom) are expressed at higher levels in pseudopalisading zones of human GBM. **(h)** IHC images and quantifications show increased abundance of GLUT1^+^ and Iba1^+^ areas in GBM of MES subtype as compared PN or CL subtypes. n=5 zones for each subtype, one-way ANOVA, \**P*<0.05, \*\**P*<0.01, \*\*\**P*<0.001, ns, not significant.

The UnaG^+^ cells were also relatively quiescent, as indicated by a downregulation of cell cycle genes (*Rgcc* and *Cdc20*), and a smaller fraction expressing the proliferation marker Ki67 (∼6% of UnaG^+^ vs. ∼30% of UnaG^-^ cells) (**Fig. 6b; Fig. S8c**). This resonates with our recent report linking GBM quiescence and HIF signaling (Tejero et al., 2019).

The top upregulated DEGs in UnaG^+^ GBM cells included many canonical HIF target genes that mediate metabolic adaptation, e.g., *Aldoa, Ldha, Slc2a1* (Glut1)*, Pgam1, Pkm, Pgk1, Gpi1, Pfkl,* etc., further confirming faithful reporting of HIF activity by HRE-UnaG. UnaG^+^ tumor cells also upregulated genes regulating angiogenesis (*Vegfa*), autophagy (*Bnip3*), apoptosis (*Eno1*), and notably, macrophage chemotaxis (macrophage migration inhibitor factor (*Mif)*) and inflammatory cytokines such as *Cxcl10* and *Cxcl9* (**Fig 6b; Table S1**). In agreement, UMAP plot showed that UnaG^+^ cells expressed higher levels of *Mif* and *Lgals3*, both known to regulate macrophage trafficking and immune suppression (Bach et al., 2009; Dumic et al., 2006; Kouo et al., 2015; Mittelbronn et al., 2011) (**Fig. S8d**).

Consistent with an engagement of immune signaling, pathway enrichment analyses by ENRICHR, Ingenuity Pathway Analysis (IPA), and Gene Set Enrichment Analysis (GSEA) (Krämer et al., 2014; Subramanian et al., 2005; Xie et al., 2021) revealed that aside from hypoxia, glycolysis, gluconeogenesis, cholesterol biosynthesis and metabolism, and Robo4-VEGF crosstalk, UnaG^+^ GBM cells are negatively enriched for Type II IFN signaling, TNF⍺ NF-κB signaling pathway, Neuroinflammation, signaling pathway, as well as IFN⍺ and IFNγ responses (**Fig. 6c; Fig. S8e, f; Fig. S9a, b**; **Table S2**). In addition, the gene ontology (GO) Negative Regulation of Immune System Process was predominantly upregulated in UnaG^+^ tumor cells, and conversely, the GO Positive Regulation of Immune System Process was downregulated in UnaG^+^ cells (**Fig. S9c**). Hence, the in vivo GBM-hypoxia gene signature underscores a high degree of immune interactions favoring immunosuppression.

### UnaG^+^ gene signature is enriched in recurrent GBM and predicts poor prognosis for GBM patients

We next compared the in vivo GBM-hypoxia gene signature captured by HRE-UnaG (top 200 upregulated DEGs) to the ‘Hallmark’ hypoxia gene set of the MSigDB collection (list of 200 genes compiled from multiple hypoxia studies, including many in vitro studies) (Liberzon et al., 2015). This revealed an overlap of 58 common genes related to glycolysis, Robo4-VEGF signaling, and oxidative stress (**Fig. S9d; Table S3**). Unique to the UnaG^+^ GBM gene set were 142 genes concerning cholesterol metabolism, actin cytoskeleton, peroxisome proliferator-activated receptor signaling (PPAR, known to regulate lipid metabolism), and notably chemokine signaling (**Fig. S9d**).

To further explore the prognostic value of the UnaG^+^ gene signature, we applied this signature to GBM patients of the TCGA database (**Fig. S10a, b**). We found that GBM patients can be stratified into two cohorts based on high or low UnaG^+^ gene signature scores, correlating to worse or better survival, respectively (**Fig. 6d**; **Table S4**). This correlation applied to all three GBM subtypes (**Fig. S10c**). The UnaG^+^ gene signature is also more represented in recurrent GBM tumors than in primary GBM, underscoring the importance of hypoxia-induced gene expression changes in GBM relapse (**Fig. 6e**).

For additional validation of the clinical relevance of the in vivo GBM and TAM-hypoxia gene signatures, we analyzed human GBM patient data from the Ivy Glioblastoma Atlas Project (Ivy GAP), which encompass gene expression profiles from distinct topographical areas of GBM, including perinecrotic zone (Pz), pseudopalisading cells (Pc), cellular tumor, leading edge, and microvascular proliferation area (Puchalski et al., 2018). Results confirmed that the UnaG^+^ gene signature was enriched in Pz and Pc relative to other areas (**Fig. 6f**; **Table S5**). Notably, the Pz and Pc showed higher expression of genes or niche factors not only expressed by hypoxic tumor cells (e.g., *MIF* and *LGALS3*), but also by entrapped hypoxic TAMs (e.g., *CD68, MRC1, CCL8*, and *IL1B*) (**Fig. 6g**). Moreover, survey of the mRNA in situ hybridization data from Ivy GAP further verified high expression levels of UnaG^+^ DEGs in pseudopalisading patterns in the peri-necrotic zone (**Fig. S10d**).

Remarkably, scRNA-seq data showed that UnaG^+^ GBM cells were not uniform, encompassing 4 subclusters, each with functional specialization based on transcriptomics (**Fig. S11a**). For instance, sc-a UnaG^+^ cells are engaged in cell cycle regulation, apoptotic signaling, and neural crest differentiation, sc-b in angiogenesis, sc-c in unfolded protein response (UPR) and focal-adhesion, and sc-d in IFN signaling (**Fig. S11b, c**). Echoing our in vivo results on a high degree of immune signaling, various UnaG^+^ subclusters were also enriched for distinct immune pathways, e.g., sc-a for cytoplasmic retention of NF-κB; sc-b for macrophage chemotaxis; and sc-d for type I and II IFN signaling. Of note, NF-κB is a transcription factor controlling numerous pro-inflammatory cytokines such as TNF, IL-1, IL-6, and IL-12 (Oeckinghaus and Ghosh, 2009), whereas cytoplasmic retention of NF-κB signifies immune repression (Lawrence, 2009), again linking tumor hypoxia and immunosuppression.

Comparative IPA similarly revealed that sc-b UnaG^+^ cells were enriched for IL6, VEGF and CXCR4 signaling, and sc-d for IFN, neuroinflammation, and necroptosis pathways (**Fig. S11d**). Consistently, IFN, interferon regulator factors (IRF) families, and IL-1β are predicted by IPA to be upstream regulators of the gene programs for sc-b and sc-d UnaG^+^ cells (**Fig. S11e**). Resonating with above results on IL-1β being released by TAMs in response to cues from hypoxic GL261 cells and its role in shaping TAM spatial patterning and maturation of pseudopalisades, the prediction that IL-1β serves as an upstream regulator for UnaG^+^ cells underscores a reciprocal signaling between hypoxic GBM cells and entrapped TAMs.

The mesenchymal (MES) GBM subtype is known for an increased TAM compartment and worse survival when compared to proneural (PN) and classical (CL) subtypes (Verhaak et al., 2010; Wang et al., 2017). We analyzed by immunohistochemistry patient GBM samples from different subtypes for the expression of IBA1 and GLUT1. Indeed, MES subtype had the highest abundance of IBA1^+^ TAMs, and this was associated with more prominent GLUT1^+^ pseudopalisades (**Fig. 6h**). In agreement, MES subtype of the TCGA GBM patient database displayed the highest enrichment for UnaG^+^ gene signature (**Fig. S12a**). The MES GBM patients also showed highest expression of the two top UnaG^+^ DEGs *MIF* and *LGALS3*, with *LGALS3* expression also being correlated with worse outcome (**Fig. S12b**). Additionally, TCGA data showed that genes or niche factors associated with hypoxic TAMs – *CD68*, *MRC1*, *CCL8* and *IL1B,* as well as CCL8 receptors *CCR1* and *CCR5–*, are all expressed at higher levels in GBM of MES subtype and predict worse outcome (**Fig. S12c**).

### Targeting hypoxic niches in GBM disrupts TAM spatial patterning and achieves better tumor control

Lastly, to further delineate a causal link between hypoxic niches and TAM patterning, we examined how targeting hypoxic cells may impact the immune landscape in GBM. To this end, we tested evofosfamide (Evo), a chemotherapy pro-drug that is activated under hypoxia (Brenner et al., 2018), and radiation therapy (RT), which is thought to increase tumor hypoxia by damaging radiosensitive blood vessels (Barker et al., 2015). Two weeks after intracranial transplantation of GL261 into B6-WT mice, animals received either sham, RT, Evo, or combined treatment for 2 weeks (**Fig. 7a**). Examination of tumor tissues after the last treatment revealed that both Evo and RT reduced the overall tumor burden, while combined treatment achieved superior tumor reduction (**Fig. 7a, b**). Regarding hypoxic cells, Evo treatment reduced but did not fully eradicate the UnaG^+^ population, while RT alone or in combination with Evo led to a near complete eradiation of UnaG^+^ cells (**Fig. 7b**).

**Figure 7.**
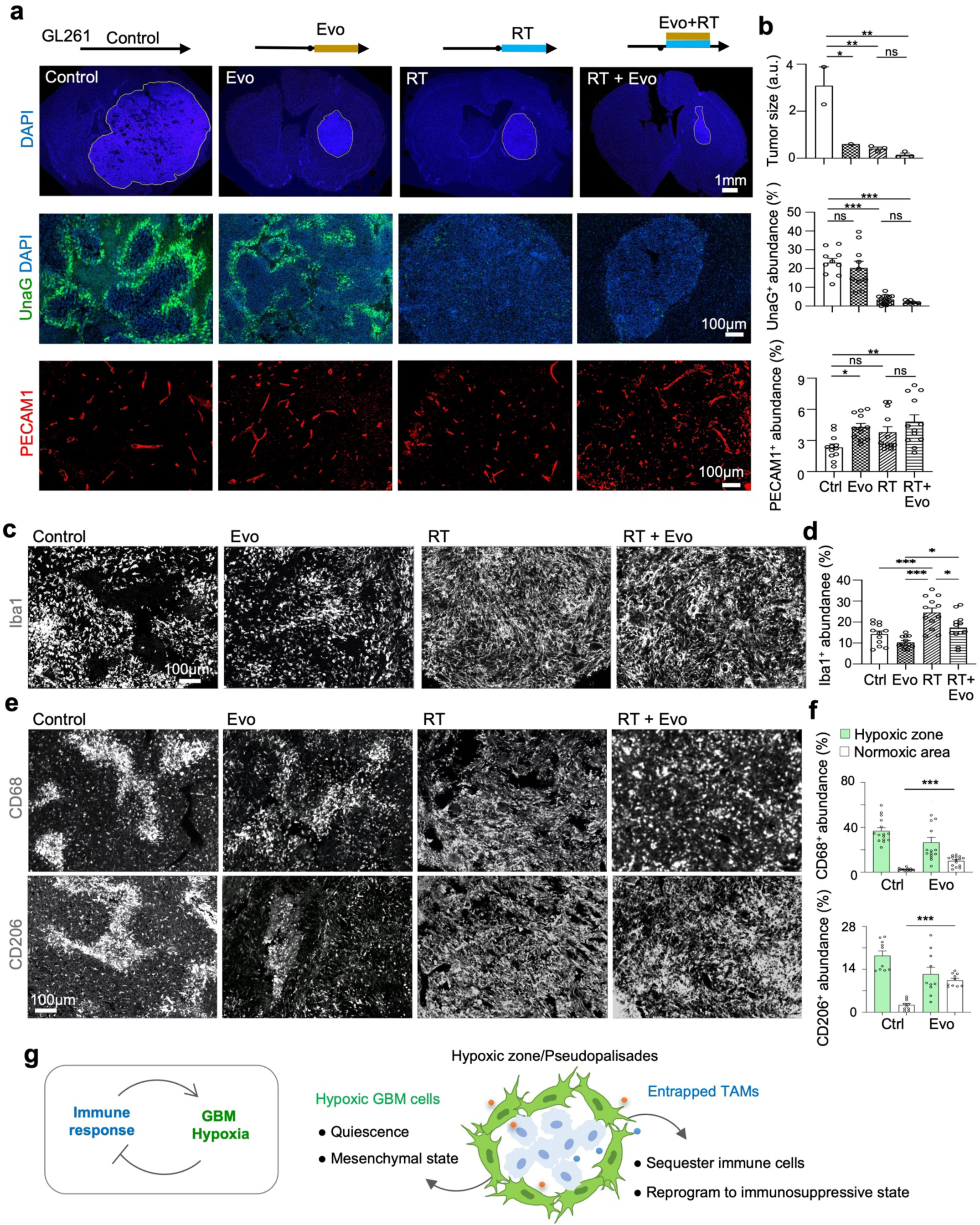
Targeting hypoxic niches in GBM disrupts TAM spatial patterning and achieves better tumor control. **(a, b)** Top, schematics: B6-WT mice bearing GL261 GBM were treated for 2 weeks with Evo, XRT, or both. Bottom, IF images and DAPI nuclear staining show that combined treatment achieved best tumor control and near eradication of hypoxic zones (UnaG^+^) as compared to treatment with Evo alone. Also note higher tumor vessel density after Evo or combined treatment. n=3 mice per cohort, one-way ANOVA, \**P*<0.05, \*\**P*<0.01, \*\*\**P*<0.001, ns, not significant. **(c, d)** IF images show higher abundance of Iba1^+^ TAMs after XRT or combined treatment. Quantification on randomly selected images from n=3 mice per treatment cohort, one-way ANOVA, \**P*<0.05, \*\*\**P*<0.001. **(e, f)** IF images show that XRT or combined treatment resulted in dissipation of of patterning of TAMs (CD68^+^ or CD206^+^) in contrast to TAM entrapment and pseudopalisades seen in control GL261 GBM. Quantification of randomly selected images from n=3 mice per group, One-way ANOVA, \*\*\**P*<0.001. **(g)** Left, diagram depicting mutual influence of immune response and tumor hypoxia – antitumor immunity exacerbates hypoxia, while hypoxia induces immunosuppression. Right, schematics of the role of hypoxia niche factors in instructing transcriptional responses of hypoxic tumor cells which exhibit quiescence and mesenchymal shift, and in promoting sequestration and reprograming of TAMs, contributing to immunosuppressive microenvironment.

It is noteworthy that HRE-UnaG reports the acute state of HIF^ON^ cells, so the abundance of UnaG^+^ cells in a particular condition reflects on-going neoangiogenesis and thereby oxygen supply in response to treatment. We assessed vascular density after different treatments, which appeared comparable in post-Evo or post-RT GBM tissues, although Evo or combined treatment results in a slightly higher vessel density (**Fig. 7a, b**). These data together suggest a higher sensitivity of hypoxic tumor cells to RT than to Evo, given the slightly lower vascular density in the former.

We next examined the impact of RT or Evo on the abundance and, importantly, the spatial patterning of TAMs. We found that in Evo-treated GBM tissues, pseudopalisading patterns were less well-defined, with areas of discontinuous UnaG^+^ cells and TAMs (Iba1^+^, CD68^+^, or CD206^+^) (**Fig. 7c-f**). In post-RT or post-RT+Evo GBM tissues, with near eradication of UnaG^+^ cells and disappearance of pseudopalisades, we found that TAMs displayed a uniform spatial distribution without confinement in specific niches; additionally, RT or combined treatment resulted in increased abundance of Iba1^+^ TAMs (**Fig. 7c-f**).

## DISCUSSION

Our study highlights hypoxic zones as a driving force in TAM patterning and immunosuppression during GBM progression. We also discovered a reciprocal impact of anti-cancer immunity on tumor hypoxia, with immune response exacerbating vascular aberrancy and thereby tumor hypoxia, while hypoxic zones attract and sequester activated immune cells and induces immunosuppression (**Fig. 7g**). The mutual influence of immune response and tumor hypoxia has important implications for improving immunotherapy against GBM.

Even though GBM is well known for its immunosuppressive TME and tumor hypoxia is linked to poor prognosis (Barsoum et al., 2014), the underlying mechanisms linking the two are not fully understood. Our studies in a fully immunocompetent model of GBM revealed that in GBM interior, majority of TAMs are blood-borne MDMs, whereas microglia largely occupied GBM periphery. Consistently, in early GBM, MDM-derived TAMs were found mostly adjacent to dense and regular blood vessels; later, vasculature became sparse, tortuous and engorged, while TAMs started to shift from perivascular regions to poorly vascularized areas. Interestingly, we found that entrapped TAMs in pseudopalisading regions also experience hypoxia, and they express immunotolerance markers such as CD206 and Arg1, as well as phagocytic marker CD68. Similarly, CTLs were also attracted to hypoxic zones, and they express T cell exhaustion markers. Hence, two potential mechanisms may account for hypoxia-induced immunosuppression: i) immune cells are attracted to and subsequently sequestered in hypoxic zones, and ii) entrapped immune cells are reprogrammed to an immunosuppressive state. The development of pseudopalisading structures in GBM thus facilitates inflammatory containment by confining cytotoxic immune cells and necrotic tumor cells (which are immunogenic) inside hypoxic cores, thereby limiting communication of antigen-presenting cells with effector immune cells.

We also discovered that the immune response has a profound effect on tumor vasculature development and thereby tumor hypoxia. Tumor hypoxia is generally assumed to arise from rapid tumor expansion outstripping blood supply; interestingly, our data unveiled immunocompetence (adaptive immunity) as an important driving force in shaping tumor vascular development and thereby tumor hypoxia. In immunodeficient SCID hosts lacking functional T and B cells, even though GBM tumors reached advanced sizes, tumor vasculature remained regular and dense with scattered hypoxic microregions that never transformed into pseudopalisades. Hence, tumor size alone does not dictate the emergence of hypoxia. In contrast, in GBM developing in immunocompetent hosts, even though the initial tumor vasculatures appeared regular and dense, they later became engorged, tortuous, and sparse. Hence, immunity-driven aberrant vascular changes may serve as major force shaping tumor hypoxia and TAM patterning.

Mechanistically, we showed that TAM trafficking and sequestration in hypoxic zones is an active process, requiring cell-cell sorting and Ccl8 and IL-1β, two cytokines expressed by TAMs in response to cues from hypoxic tumor cells. IL-1β is also identified as an upstream regulator that shapes the transcriptional responses of neighboring hypoxic GBM cells. Similar scenarios of reciprocal communication via cytokines have been also described between the predicted cancer stem cell (CSC) population of GBM and infiltrating immune cell populations (Bayik and Lathia, 2021), and it will be interesting for future studies to examine phenotypic alignment of hypoxic GBM cells and CSCs.

In this context, it should be noted that the in vivo HIF^ON^ gene signature that we revealed with the sensitive HRE-UnaG reporter (Erapaneedi *et al*., 2016) indicated that hypoxic GBM cells are equipped for metabolic adaptation and stress response and also display features of quiescence and MES, both linked to malignant potency (Bhat et al., 2013; Tejero *et al*., 2019). The UnaG^+^ GBM signature has clinical prognostic value: it is more represented in recurrent GBM, in MES-subtype and predicts shorter survival.

We also confirmed that hypoxic niche genes associated either with hypoxic GBM cells (*MIF, LGALS3*) or hypoxic TAMs (*CCL8, IL1B, CD68, MRC1*) are expressed at higher levels in perinecrotic zones or pseudopalisading cells in GBM patients, particularly of the MES subtype, which is linked to high malignant potency and enhanced immune response (Wang *et al*., 2017). Our results thus substantiate a recent report on tumor and immune cell interactions in driving the transitions to MES-like states in GBM patients (Hara et al., 2021).

From a therapeutic point of view, we showed that targeting the hypoxic niche may synergize with conventional chemoradiation treatment, by not only reducing the hypoxic tumor cells (which are therapy-resistant as chemodrugs target mainly proliferative cells, while RT requires oxygenation to be effective), but likely also by attenuating immunosuppression. Indeed, we found that combining Evo and RT eradicated hypoxic niches and TAM sequestration, and achieved better tumor control. Current clinical trials with evofosfamide for recurrent GBM have so far yielded only limited positive results for patients (Brenner et al., 2021), but future additional combinatorial strategies may be better equipped to bring out therapeutic benefit from perturbing hypoxic niches in GBM. Our results also raise the awareness that while boosting anti-cancer immunity is desirable, it may lead to aberrant tumor vasculatures, higher hypoxic burden, and immunosuppression, leading to dampened efficacy of immunotherapy.

In summary, the mutual communication between tumor and immune cells in the hypoxic niche plays a determining role in sculpting the immune landscape, limiting inflammatory spread and inducing an overall tolerogenic/immunosuppressive microenvironment. This new understanding of the reciprocal influence of immune response and tumor hypoxia will have important clinical ramifications for prognosis and development of immunotherapy for GBM patients.

## METHODS

### GBM cell lines

The murine high-grade glioma cell line GL261 (*Kras*^G12T^, *Trp53*^G153C^, *Pten*^-/-^, *c-myc* and *Egfr* amplification) (Newcomb and Zagzag, 2009; Oh et al., 2014; Szatmári et al., 2006) was obtained from the repository of the National Cancer Institute. The human patient-derived GBM cell line GBM2 was obtained from the laboratory of Dr. Foty at Rutgers University (Carminucci et al., 2020). Glioma cells were cultured in DMEM media with Glutamax (Gibco) supplemented with 10% FBS (Thermo Fisher Scientific) and 1% Penicillin-Streptomycin antibiotics (Gibco). For long term storage, cells were cryopreserved in medium containing 10% DMSO. Cells were passaged after thawing for at least two passages before use in experiments. For in vitro hypoxia studies, GL261 cells were seeded into 6 well plates and placed for 24 hr in a hypoxia chamber (C-Chamber; Biospherix) set to 1% oxygen.

### Generation of lentiviral HIF reporter

The plasmid dUnaG was obtained from the Kiefer laboratory (Erapaneedi *et al*., 2016) and used as template to amplify an HRE-dUnaG PCR fragment, which was inserted into the Gateway entry vector pENTR/D-TOPO (Invitrogen) and then transferred by Gateway LR reaction (Invitrogen) into the destination plasmid pLenti X1 Puro DEST (Addgene #17297) (Campeau et al., 2009). The final pLenti-HRE-dUnaG plasmid (deposited as Addgene #124372) carries an array of five hypoxia-response elements (HRE) and a minimal CMV promoter to drive the expression of UnaG fused to a PEST degron and C-terminal myc tag, and in addition a puromycin resistance gene driven by a constitutive phosphoglycerate kinase (PGK) promoter. Of note, the myc tag was used in some immunofluorescence experiments to stain for UnaG with an anti-myc antibody to extended stability of fluorescence for storage of sections.

Lentiviral particles were produced by transfection of HEK293T cells with the pLenti plasmid, envelope plasmid pMD2.G, and packaging plasmid psPAX2 (Addgene #12259 and #12260; deposited by Didier Trono, EPFL Lausanne). Media supernatants were collected 2–3 days after transfection and viral particles were concentrated by ultracentrifugation.

### Transduction of GBM cells

GL261 cells were transduced with a mixture of HRE-UnaG lentiviral particles and polybrene (5 mg/ml). Transduced cells were selected with 1 µg/ml puromycin, starting 48 hr after transduction. Early passages of GL261-HRE-UnaG cells were frozen 1 wk after puromycin selection, and subsequent passages were cultured in media with 0.5 µg/ml puromycin to prevent silencing of lentiviral vectors.

### Mice

C57BL/6J wild-type mice and C57BL/6J-SCID mice were obtained from The Jackson Laboratory. ICR-SCID mice were purchased from Taconic Biosciences. *Ccl8/12* KO mice were gifted by Dr. Sabina Islam (*45*) and *Il1b* KO mice were gifted by Dr. Dmitry Shayakhmetov (Horai et al., 1998), and were both bred in our colony on C57BL/6J genetic background. Mice of both genders in the age range of 8-16 weeks were used for experiments. All animals were housed in a climate-controlled, pathogen-free facility with access to food and water ad libitum under a 12-hour light/dark cycle. All animal procedures were conducted in accordance with protocols approved by the Institutional Animal Care and Use Committee (IACUC) of Icahn School of Medicine at Mount Sinai.

### Intracranial transplantation of tumor cells

Mice were anaesthetized in an induction chamber with a 2.5% isoflurane/oxygen mixture and secured to a stereotaxic apparatus (Stoelting). Anesthesia was maintained with a 1.5% isoflurane/oxygen mixture, which was delivered via a nose-cone. A lubricant ophthalmic ointment (Artificial Tears, Akron) was applied. A cranial hole was drilled through a scalp incision 2.0 mm lateral and 0.5 mm posterior to Bregma. GBM cells (10^5^ cells) suspended in 5 µl PBS were then injected through the hole at a depth of 3.2 mm using a 10 µl gas tight syringe (Hamilton) and a Nanomite programmable syringe pump (Harvard Apparatus) with a constant infusion rate of 1 µl/min to prevent backflow. After injection, syringe was incrementally raised using the stereotaxic apparatus over a period of 5 min. Scalp incision was sealed using a tissue adhesive (Vetbond, 3M).

### Immunofluorescence and immunohistochemistry

Mice carrying intracranial GBM transplants or *de novo* tumors induced by RCAS/t-va system were intracardially perfused with PBS followed by 4% PFA/PBS. Brains were harvested and fixed overnight in 4% PFA/PBS followed by two successive overnight incubations in 12.5% and 25% sucrose/PBS. Brains were then embedded in O.C.T compound (Fisher Scientific), frozen on dry ice. Cryosections of 20 µm thickness were cut using a cryostat (Leica) and collected in PBS and stored as floating sections at 4°C.

For immunofluorescence, floating sections were blocked for 1 hr (blocking buffer: PBS with 5% donkey serum and 0.3% Triton X-100), then incubated overnight with primary antibodies in antibody dilution buffer (PBS with 1% BSA and 0.3% Triton X-100), followed by staining with Alexa-labeled secondary antibodies (Jackson ImmunoResearch) for 2 hr, and counterstaining with DAPI (Invitrogen). Sections were washed in PBS and mounted with Fluoromount G (Southern Biotech).

For immunohistochemistry of microtome sections from patient GBM specimens and from mouse brains carrying RCAS induced tumors, sections from FFPE tissue blocks were processed at the Mount Sinai Pathology core using a Ventana system (Roche).

The following primary antibodies were used:

anti-Arg1 (host species: goat), Santa Cruz Biotechnology sc-18355, 1:100 dilution;
anti-CD68 (rat), Bio Rad MCA1957GA, 1:100;
anti-CD8a (rat), eBioscience 14-0081-82, 1:100;
anti-CD206 (goat), R&D Systems AF2535, 1:100;
anti-CD16/32 (rat), BD Biosciences 553142, 1:100;
anti-Foxp3 (rat), eBioscience 14-5773-82, 1:100;
anti-GLUT1 (rabbit), eBioscience MA5-31960, 1:200;
anti-Iba1 (rabbit), Fujifilm Wako 019-19741, 1:500;
anti-Integrin alpha4 (rat), BioLegend 103701, 1:100;
anti-Ki67 (rabbit), Abcam ab15580, 1:200;
anti-myc-tag (goat), Novus Biologicals NB600-335, 1:200;
anti-PECAM1 (rabbit), Abcam ab28364, 1:50;
anti-PU.1 (rabbit), eBioscience MA5-15064, 1:200.

### Single-cell RNA sequencing

Animals were euthanized, brain tissue containing the main tumor bulk was dissected on ice, and a tissue piece of approximately 3 mm edge length was diced with scalpel blades and dissociated into single cell suspension using papain digestion (Miltenyi Neural Tissue Dissociation Kit (P) 130-092-628).

The cell suspension was pelleted and resuspended in 7 ml of HBSS (without Ca/Mg), mixed with 1.2 ml of fetal bovine serum (FBS) and 3.6 ml of 100% Percoll (GE Healthcare). The Percoll cell suspension was overlaid with 1 ml of 10% FBS in DMEM and spun at 800g for 15 min, and pellet was collected in a new 15 ml tube and resuspended in 0.5 ml of FACS buffer ((Hibernate-E low fluorescence (BrainBits) with 0.2% BSA and 20 μg/ml DNase (Worthington)). Red blood cells (RBCs) were lysed by incubating cells with RBC lysis buffer (BioLegend) for 15 min at room temperature; cells were washed and resuspended in FACS buffer. The final cell suspension was submitted for single cell sequencing with the 10X Genomics system at the Mount Sinai Genomics core facility (using ∼10,000 viable cells from the sample).

### Bioinformatic analysis

Clustering analysis of scRNA-seq data was performed with the Seurat software package on the R platform (Stuart et al., 2019). We calculated signature score for GBM cell state with the scrabble software package (jlaffy.github.io/scrabble/), using the gene list of the MES2 signature (Neftel *et al*., 2019).

Gene set enrichment analysis (GSEA) of gene lists ranked by expression changes was performed with the GSEA platform (Mootha et al., 2003; Subramanian *et al*., 2005). We used the ENRICHR website for the pathway analysis for differentially expressed genes (cut-off: *P* < 0.05; fold-change > 2-fold) (Xie *et al*., 2021). The GBM biodiscovery portal (http://gbm-biodp.nci.nih.gov; accessed 02/2021) was used to match UnaG^+^ gene expression signature with patient survival (Celiku et al., 2014). cBioPortal (https://www.cbioportal.org; accessed 02/2021) was used to match occurrence in recurrent vs. primary GBM (TCGA-GBM Firehose legacy dataset, mRNA expression (mRNA expression z-scores relative to all samples (log RNA Seq V2 RSEM)) (Cerami et al., 2012; Gao et al., 2013). Pseudotime analysis was performed with Monocle 3 software package on R platform (Cao et al., 2019). The Ivy GAP database, containing expression data from 122 micro-dissected anatomical domains of 10 GBM patients, was used for GBM anatomical transcriptional analysis (glioblastoma.alleninstitute.org) (Puchalski *et al*., 2018).

### RCAS glioma samples

A genetically modified mouse model using the RCAS/t-va system was used to generate murine GBM, as previously described (Herting *et al*., 2019). Briefly, DF1 avian fibroblast cells (ATCC) were grown at 39°C, expanded to passage 4 and transfected with RCAS-PDGFB-HA or RCAS-shp53-RFP using a Fugene 6 Transfection kit (Roche, 11814443001). Cells were cultured with DMEM media (Gibco, 11995-065) supplemented with 1% L-glutamine, 1% penicillin/streptomycin, and 10% FBS (ATCC). Transfected DF1 cells were used for injections before reaching passage 25. DF-1 cells (4x10^4^) were stereotactically delivered with a Hamilton syringe equipped with a 30-gauge needle for tumor generation. The injection site was the frontal striatum with the coordinates AP -1.7 mm and right -0.5 mm from bregma; depth -2.0 mm from dural surface. Mice were continually monitored for signs of tumor burden and were sacrificed upon observation of endpoint symptoms including head tilt, lethargy, seizures, and excessive weight loss.

### Pimonidazole staining

Mice bearing intracranial GL261 tumors were intraperitoneally injected with 60 mg/kg pimonidazole (Hydroxyprobe), which was diluted in 0.9% saline. To stain for pimonidazole labeled cells, brain cryosections were blocked overnight and incubated the with mouse Dylight-549 anti-pimonidazole antibody (clone 4.3.11.3; Hydroxyprobe). Sections were mounted with Fluoromount-G (SouthernBiotech) and images were acquired by fluorescence microscopy with a Zeiss Axio microscope.

### Irradiation and evofosfamide treatments

Mice bearing GBM were randomly divided into four cohorts (Control, XRT only, Evo only, XRT+Evo). Treatments were administered two weeks after intracranial transplantations to allow for tumor establishment. For XRT, mice were irradiated in a X-Rad 320 irradiator (Precision X-Ray), twice every week with a 2.5 Gy dose for two weeks (5 Gy total dose/week). A lead shield was placed over the body of the mice to only expose heads to radiation. Mice in the Evo cohort were administered with 50 mg/kg i.p. injection of Evo (AdooQ Bioscience) every day for two weeks. Mice in the XRT+Evo treatment cohort were injected daily with Evo (50 mg/kg) for two weeks and were concurrently treated with 2.5 Gy radiation twice every week. Mice in all cohorts were weighed daily and monitored for ambulatory, feeding, and grooming activities, and animals meeting humane endpoints were euthanized.

### qRT-PCR analysis of bone marrow derived macrophages treated with tumor conditioned media

The isolation of bone marrow-derived macrophages followed previously described protocols (Bailey et al., 2020). In brief, mice were sacrificed by cervical dislocation and bone marrow was extracted from tibia and femur by flushing out with DMEM media containing 10% FBS with a 25-gauge needle attached to a 10 ml syringe. The cell suspension was passed through a 70 µm strainer and centrifuged at 300 g for 5 min at room temperature. The cell pellet was resuspended with DMEM media containing 10% FBS and MCSF (25 ng/ml; Peprotech). The media was changed every two days for 7 days to obtain pure differentiated BMDMs.

To produce conditioning media, 300,000 cells of murine GBM cell lines GL261 (Szatmári *et al*., 2006) were plated in 10 cm culture dish in DMEM media containing 10% FBS. After one day, media was changed to fresh media and the cells were incubated for another 2 days. To obtain media supernatant conditioned by hypoxic GBM cells, dishes were placed in a 1% oxygen hypoxic chamber (Biospherix). The conditioned media was collected after 2 days and the supernatant was centrifuged at 300 g for 5 min to remove cell debris.

BMDM cultures were treated with supernatant for 48 hours. For qRT-PCR analysis, RNA of BMDMs was extracted using the RNeasy mini kit (QIAGEN) and cDNA was synthesized using SuperScript III First Strand Synthesis System (Invitrogen). Quantitative PCR was performed using PerfeCTa SYBR Green FastMix Rox (Quanta Biosciences) in the ABI 7900HT qPCR system (Applied Biosystems).

The following primers were used:

Arg-1-F: GTGGCTTTAACCTTGGCTTG; Arg-1-R: CTGTCTGCTTTGCTGTGATG;
Ccl8-F: ACAATATCCAGTGCCCCATG; Ccl8-R: TGAAGGTTCAAGGCTGCAG;
CD68-F: ACTTCGGGCCATGTTTCTCT; CD68-R: GCTGGTAGGTTGATTGTCGT;
Il1b-F: CCAAGCAACGACAAAATACC; Il1b-R: GTTGAAGACAAACCGTTTTTCC;
Gapdh-F: ACTGCCACCCAGAAGACTGT, Gapdh-R: GATGCAGGGATGATGTTCT;
Gapdh was used as the housekeeping gene for normalization.

### ELISA of bone marrow derived macrophages treated with tumor conditioned media

ELISA was performed using the Mouse IL-1 beta/IL-1F2 DuoSet ELISA kit (R&D Systems DY401-05), using additional reagents from the DuoSet ELISA ancillary reagent kit 2 (R&D Systems DY008). Briefly, a 96-well microplate was coated with IL-1β capture antibody, then 100 µl of samples or standards was added to each well, followed by streptavidin-HRP, and color reaction. The plate was then immediately read at 450 nm using a microplate reader. Microsoft Excel and GraphPad Prism 9 software were then used to generate a standard curve to calculate values for samples.

### Image quantifications

Image data were quantified with Fiji Is Just ImageJ (FIJI) package of Image J (Schindelin et al., 2012). To quantify abundance of marker or UnaG reporter expression, in each selection of region of interest (ROI), percentage of ROI area positive for fluorescent or immunosignals was automatically counted after applying a threshold mask followed by morphological segmentation using MorphoLib J Plugin (Legland et al., 2016). For vessel density, in each selected region of interest (ROI), PECAM1^+^ vessel sections were manually counted and numbers were normalized to area size of ROI. To quantify signal abundance per unit area, images were spatially calibrated and total number of signal positive areas were divided by area of ROI. The ‘plot profile’ function of Image J was used to generate profiles of relative immunofluorescence in a chosen ROI (this function sums up the pixel values of vertical lines in a ROI and plots them as profile).

### Statistical analysis

Statistical analyses were performed using GraphPad Prism 9 software. Bar graphs represent means and error bars represent SEM. One-way ANOVA with Tukey’s post-hoc correction (for three or more experimental groups) and t test were performed to assess if experimental groups were significantly different from each other. *P* < 0.05 was considered to be statistically significant (*). *P* < 0.01, **; *P* < 0.001, ***.

## ACKNOWLEDGEMENTS

We thank Friedemann Kiefer (Max-Planck Institute for Molecular Biomedicine, Münster, Germany) for sharing HRE-UnaG plasmids, Kristin Beaumont (Icahn School of Medicine at Mount Sinai) for advising on single-cell RNA sequencing experiments, and Sanjana Shroff (Mount Sinai Genomics Core Facility) for performing quality control and library preparations for single-cell RNA sequencing. We also thank Ruben Fernandez-Rodriguez (Mount Sinai Biorepository and Pathology Center) for help with IHC of patient specimens; and Glenn Doherty and Nikolaos Tzavaras (Mount Sinai Microscopy Core Facility) for assisting in imaging. We thank Dalia Halawani and all members of Zou and Friedel laboratories for their valuable suggestions.

## AUTHOR CONTRIBUTIONS

Conceptualization: AS, RHF, HZ

Methodology: AS, ZC, SK, AR, LS, DH, RHF, HZ

Investigation: AS, VM, ZC, SK, CB, AR Supervision: RHF and HZ

Writing – original draft: AS, RHF, HZ

Writing – review & editing: AS, ZC, LS, DH, RHF, HZ

## DATA AND MATERIALS AVAILABILITY

All data are available in the main text or the supplementary materials. The lenti-HRE-UnaG plasmid has been deposited at Addgene. Single cell RNA-seq data has been deposited at the NIH NCBI GEO data repository (GSE179077). *Reviewer access token: ktgfeecanvctfsx*.

## SUPPLEMENTARY MATERIAL

**Figure S1.**
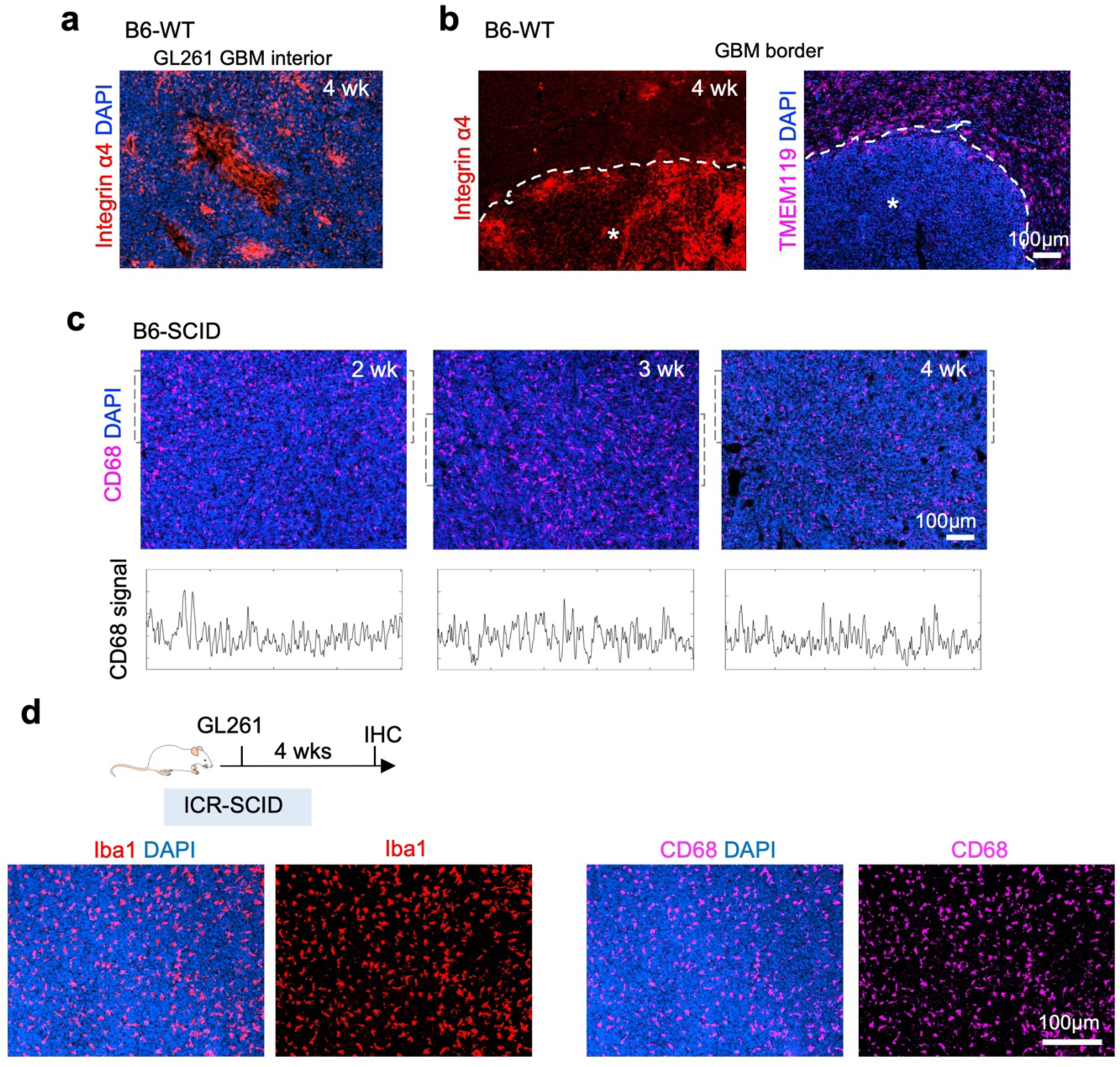
Different spatial organization of TAMs in GL261 GBM transplanted in hosts with different immunocompetence status. **(a)** IF images of GL261 tumor growing for 4 weeks in the brain of B6-WT host. Note that monocyte-derived macrophages (MDMs, Integrin α4^+^) located in tumor interior displayed a distinct spatial pattern. **(b)** IF images show tumor border of GL261 GBM denoted by dashed line. Asterisks denote tumor interior. MDM (Integrin α4^+^) were mostly located in tumor interior, while microglia-derived TAMs (TMEM119^+^) were mainly present in peritumoral regions. **(c)** IF images show that in GL261 GBM transplanted in B6-SCID mice, CD68^+^ TAMs were uniformly distributed throughout GBM progression. **(d)** GL261 GBM established in ICR-SCID host mice harbored a relatively uniform distribution of Iba1^+^ and CD68^+^ TAMs.

**Figure S2.**
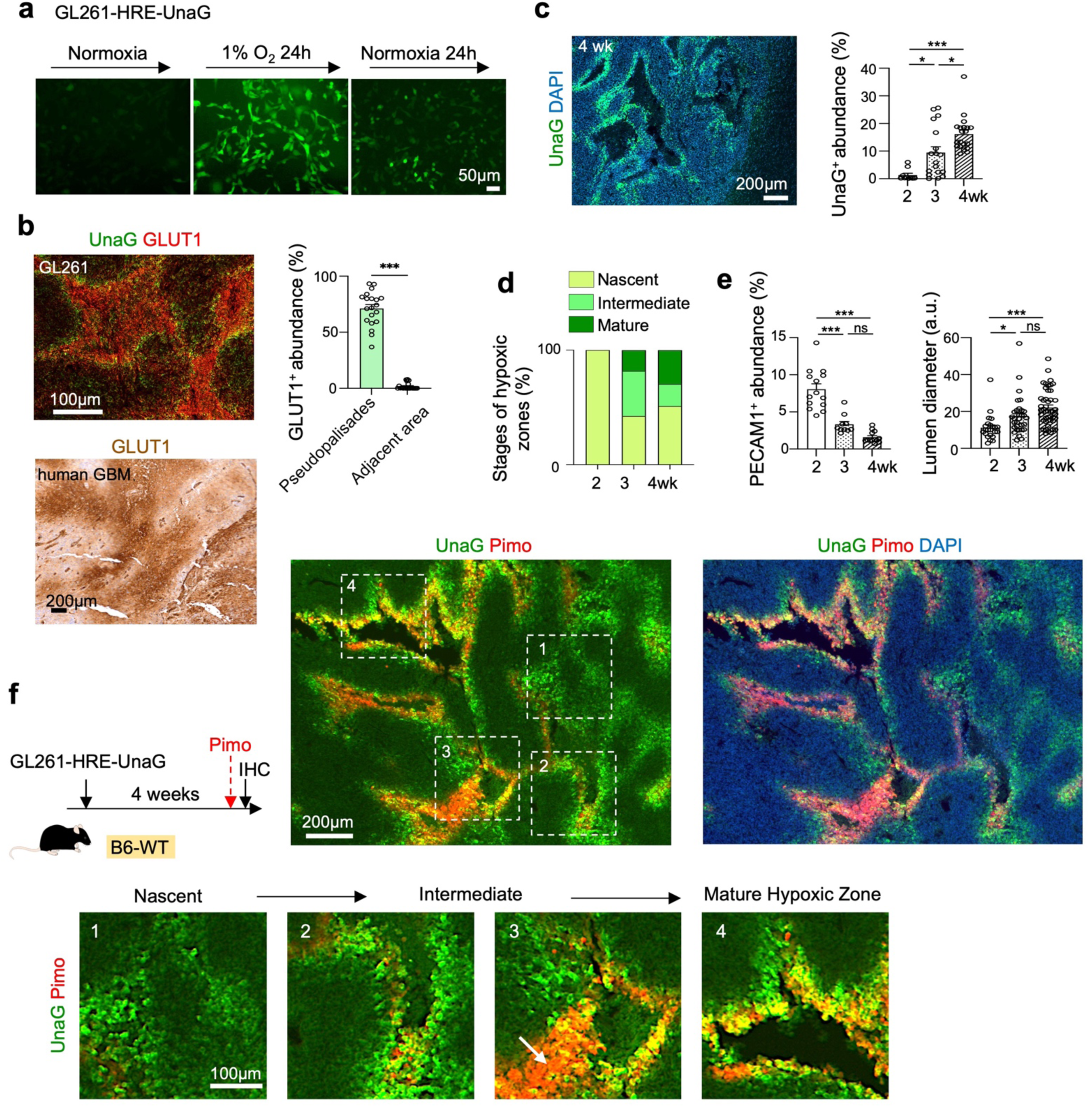
HIF reporter labels hypoxic GBM cells in GL261 GBM model. **(a)** GL261 murine GBM cells transduced with lenti-HRE-UnaG upregulated UnaG upon exposure to 1% O_2_ and turned off UnaG when returned to normoxic conditions. **(b)** Top, IF images show overlap of UnaG signals with GLUT1 (glucose transporter 1) in pseudopalisading structures, confirmed by quantification shown on right. n=4 mice per group, unpaired t-test; \*\*\**P* <0.001. Also note the presence of GLUT1^+^ stromal cells (UnaG^-^) in the center of hypoxic cores, likely representing hypoxic immune cells. Bottom, human GBM patient tissue (mesenchymal subtype), stained for GLUT1 revealed similar spatial patterning of pseudopalisades. **(c)** IF image shows representative example for UnaG^+^ pseudopalisades in GL261 GBM transplanted in B6-WT host at 4 weeks. Quantification shows increased abundance of UnaG^+^ area during tumor progression. n=3 mice per timepoint, one-way ANOVA; \**P*<0.05, \*\*\**P*<0.001, ns, not significant. **(d)** Quantification shows maturation of UnaG^+^ hypoxic zones from nascent (small UnaG^+^ aggregates) to mature pseudopalisades (UnaG^+^ cells encircling necrotic cores) during GL261 progression. **(e)** Quantifications show progressively reduced tumor vasculature density along with vessel lumen engorgement. n=3 mice per timepoint, one-way ANOVA, \**P*<0.05; \*\*\**P*<0.001; ns, not significant. a.u., arbitrary units. **(f)** Left, experimental scheme with injection of hypoxia-sensitive labeling compound pimonidazole (Pimo) 1.5 hours before tissue collection. HRE-UnaG labeled a wider field than Pimo. Enlarged images of boxed areas are shown below. Arrow in box 3 marks cells inside hypoxic core that were labeled by Pimo but not by UnaG, representing entrapped stromal cells. In mature hypoxic zones in pseudopalisading pattern (box 4), UnaG signals largely overlapped with Pimo.

**Figure S3.**
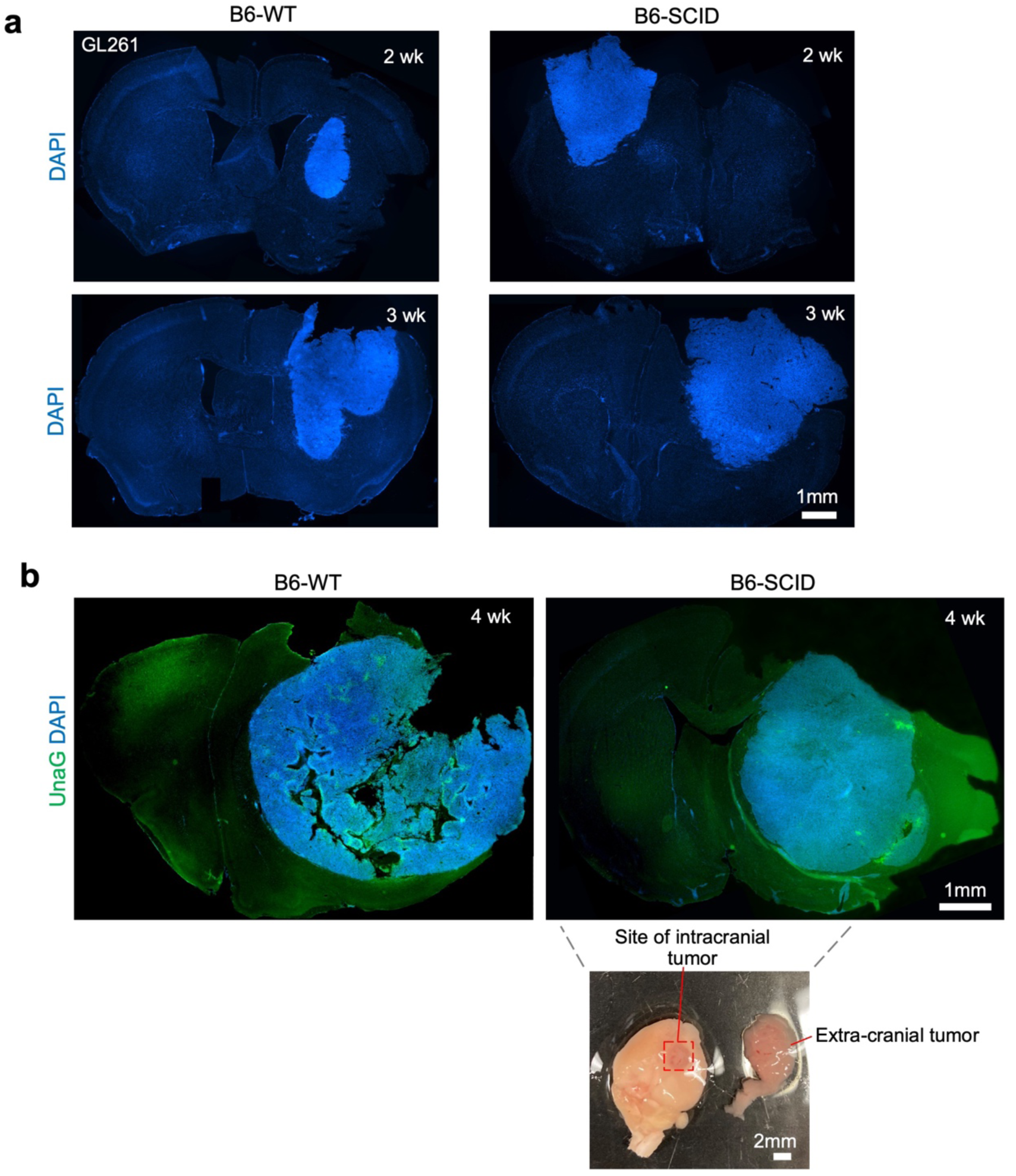
Immunocompetence of host animal impacts tumor growth and hypoxic burden. **(a)** GL261 GBM transplanted in B6-SCID hosts expanded into larger tumor mass when compared to similarly established GBM in B6-WT mice at matching time points. **(b)** GL261-HRE-UnaG GBM in B6-WT host at 4 week post-transplant contained abundant UnaG^+^ pseudopalisades, which were largely absent from GBM established in B6-SCID hosts. Note that GL261 GBM in B6-SCID mice grew aggressively and often expanded extra-cranially through the skull trephination, shown in the photo below.

**Figure S4.**
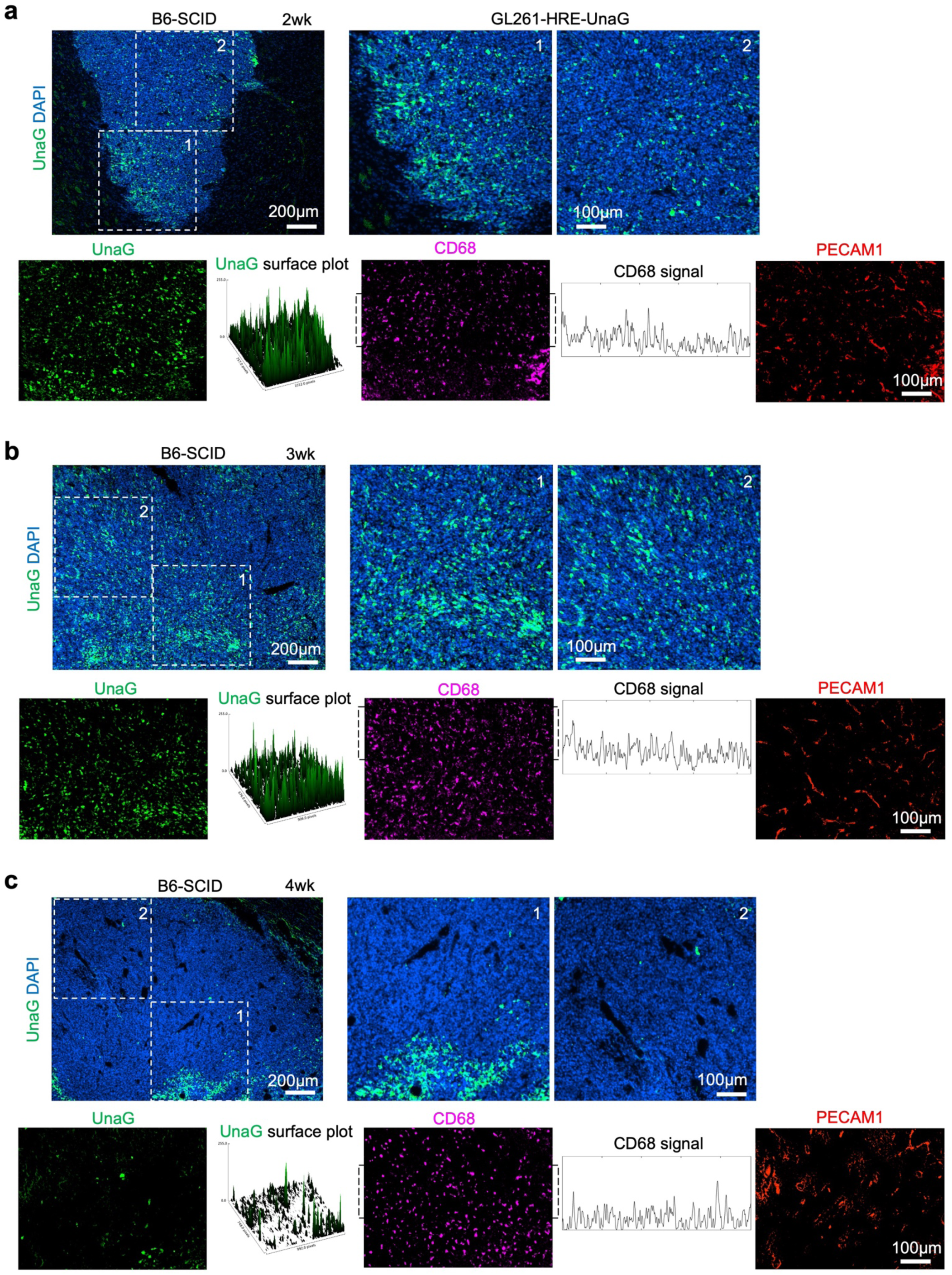
Emergence and dissipation of UnaG^+^ GBM cells during GL261 progression in B6-SCID host mice. **(a, b)** Top, IF images of GL261 GBM established in B6-SCID mice at 2 weeks (a) or 3 weeks (b) post-transplant show abundant UnaG^+^ HIF^ON^ GBM cells scattered throughout the tumor. Enlarged images of dashed boxes are shown on right. Bottom, 3D surface plots of UnaG signals (left) or relative signal intensity plot of CD68 (middle) within the regions of interest (ROI) show lack of pseudopalisading patterns paralleled by dense regular vasculature (PECAM1^+^). **(c)** Top, IF images of GL261 GBM established in B6-SCID mice at 4 weeks post-transplant show resolution of UnaG^+^ HIF^ON^ GBM cells in majority of tumor tissue (box 2), with exception of small clusters that did not form pseudopalisades (box 1). Enlarged images of dashed boxes are shown on right. Bottom, 3D surface plots of UnaG signal (left) or relative signal intensity plot of CD68 within ROI (middle) show scant UnaG^+^ cells, paralleled by uniform distribution of CD68^+^ TAMs and dense vasculature.

**Figure S5.**
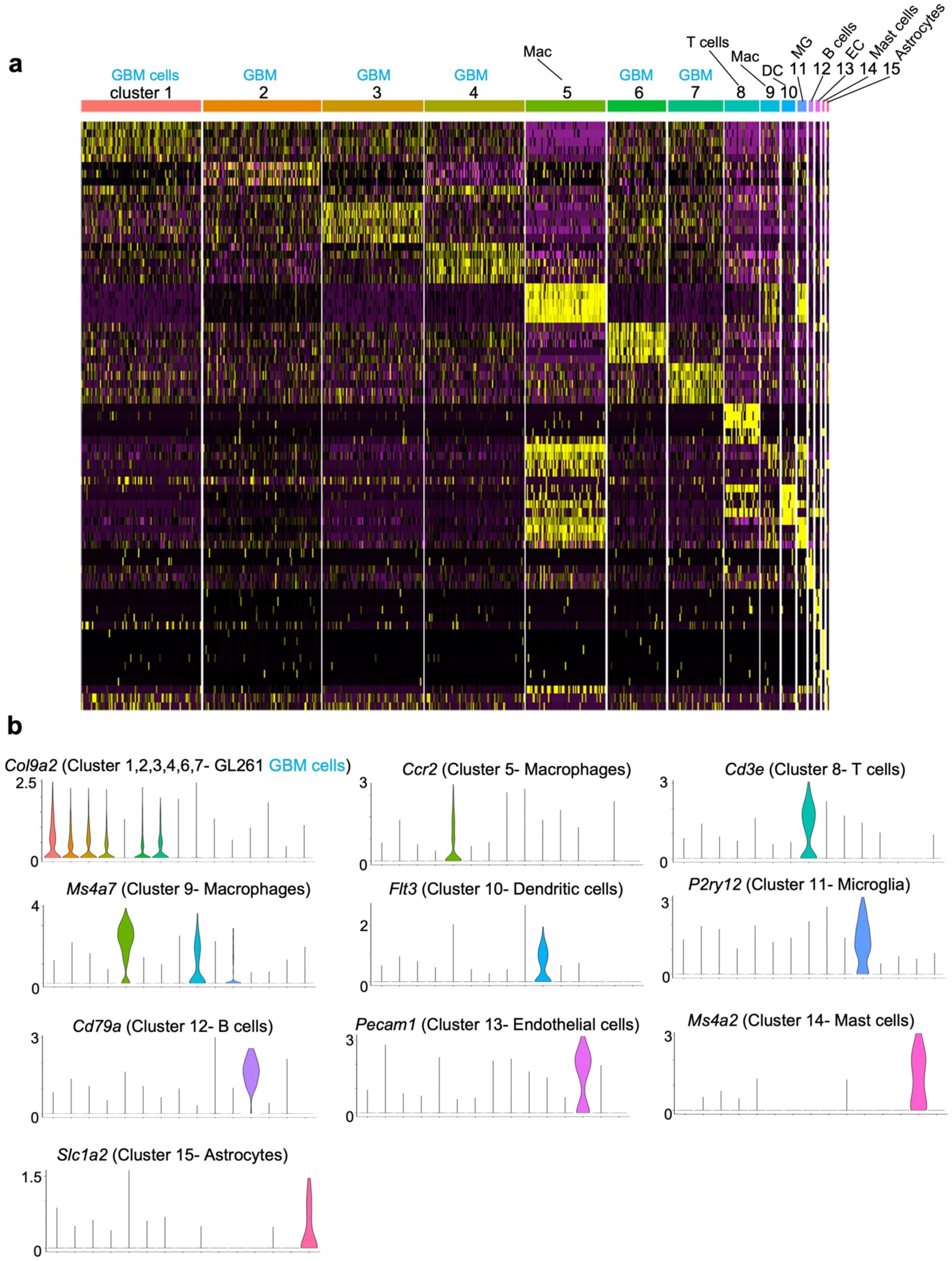
Annotation of cell types from single cell RNA sequencing of GL261 GBM. **(a)** Heatmap of top differentially expressed genes (DEGs) among all 15 cell clusters using Seurat R package. Cell types are assigned using Panglao DB online database and cell type defining gene markers. **(b)** Violin plots show unique expression of marker genes used to annotate cell types.

**Figure S6.**
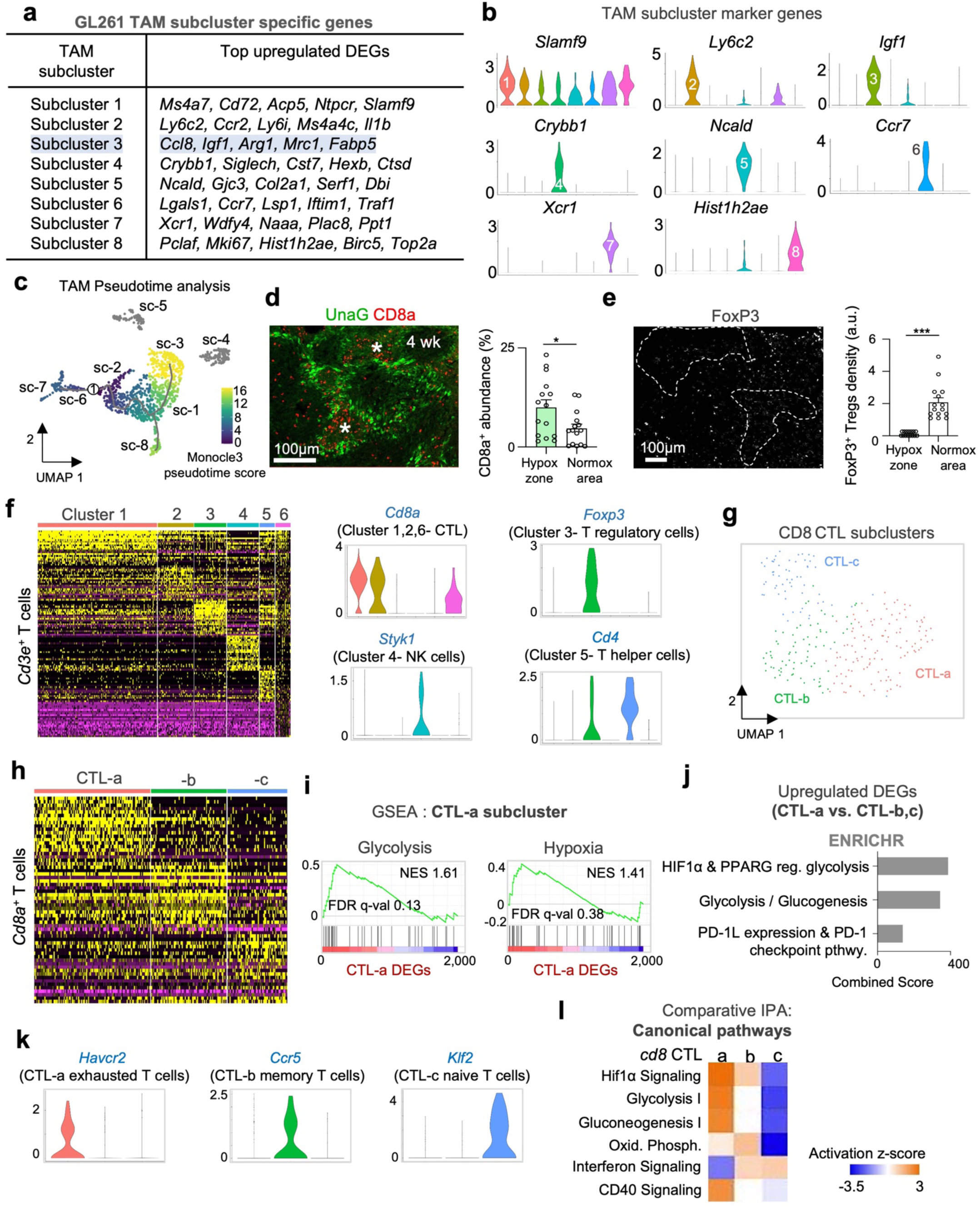
scRNA-seq analysis reveals heterogeneity of TAMs and T cells in GL261 GBM with hypoxia and immunosuppressive subpopulations. **(a)** Table of top upregulated genes in TAM subclusters. Subcluster 3 (highlighted in blue) consists of hypoxic TAMs, while subcluster 4 represents microglia. Note top upregulated DEGs in sc-3 TAMs included genes highly expressed by entrapped TAMs shown by IF images, including Arg1, Mrc1 (CD206), and CD68. **(b)** Violin plots show expression levels of selected marker genes for each TAM subcluster. **(c)** Pseudotime analysis of TAM subclusters predicts a differentiation hierarchy starting from node 1 (subcluster 2 consisting of blood-borne monocytes (*Ccr2*^+^)) with lineage branching to more differentiated clusters, including subcluster 3, which consists of hypoxic TAMs. **(d)** IF image of GL261 GBM at 4 weeks post-transplant and quantification show confinement of CD8 CTLs within hypoxic cores (asterisks). n=3 mice per group, unpaired t-test, \**P*<0.01. **(e)** IF images and quantification show that FoxP3^+^ regulatory T cells (Tregs) were distributed away from UnaG^+^ zones (dashed lines). n=3 mice per group, unpaired t-test, \*\*\**P*<0.001. **(f)** Left, heatmap shows top DEGs for T cell subclusters. Right, violin plots show selectively expressed genes used to annotate cell type for each subcluster. **(g, h)** UMAP and heatmap show CD8 CTL subclusters and the top DEGs. **(i)** Gene set enrichment analysis (GSEA) reveals glycolysis and hypoxia as top positively enriched gene sets in CD8 CTL subcluster-a. **(j)** ENRICHR analysis of the 276 genes upregulated in CTL subcluster a shows enrichment of genes involved in hypoxia, and PD-1L PD-1 checkpoint pathways. **(k)** Violin plots show marker genes for each CD8 CTL subcluster. Note that subcluster-a uniquely upregulated T cell exhaustion maker *Havcr2*. **(l)** Comparative IPA shows that subcluster-a CTLs engage in hypoxia responses, and are associated with downregulated interferon signaling.

**Figure S7.**
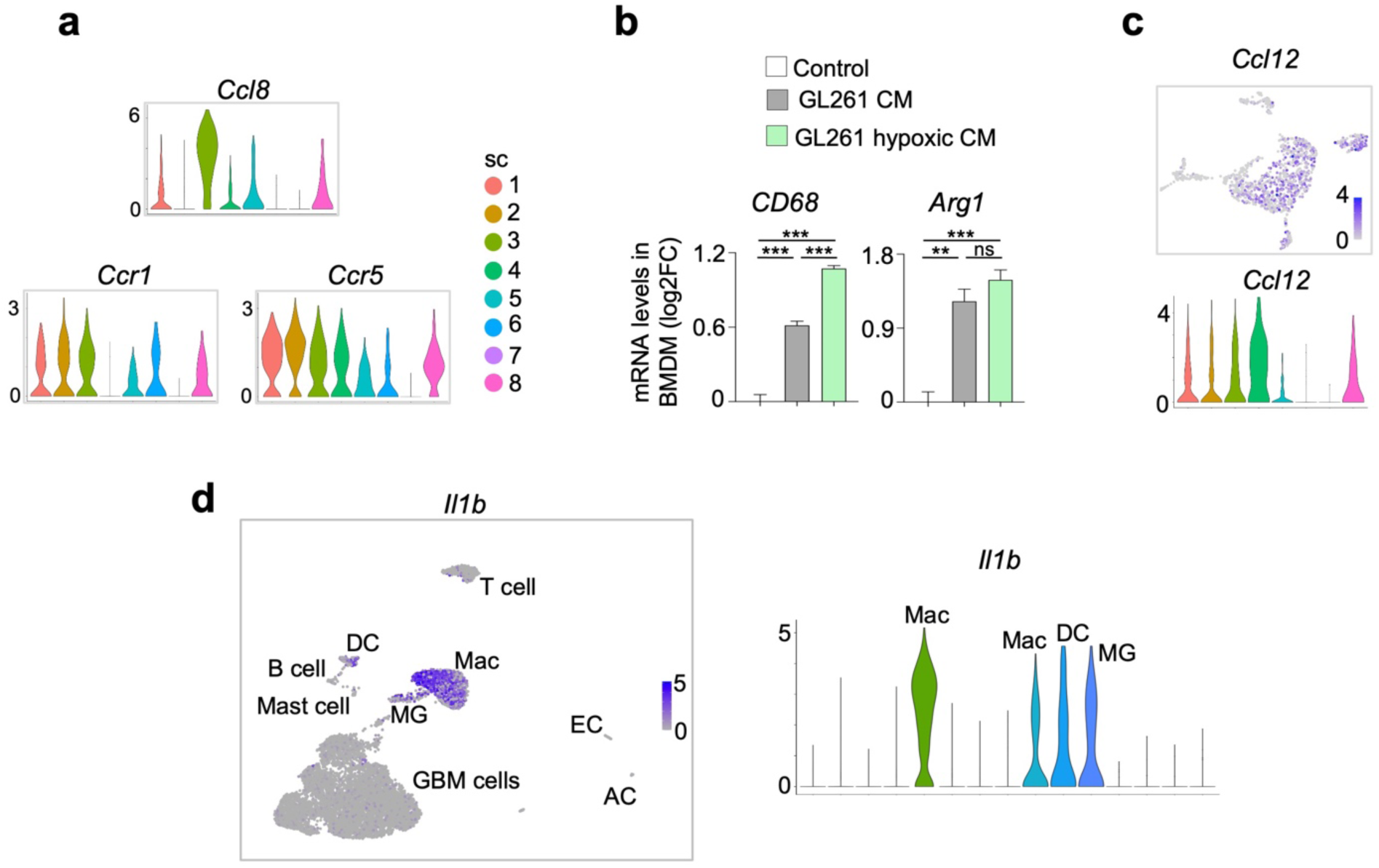
Hypoxic GBM cells release factors to induce *Ccl8* and *Il1b* expression by TAMs. **(a)** Violin plots show expression of chemokine *Ccl8* and its receptors (*Ccr1* and *Ccr5)* in TAM subclusters based on scRNA-seq data of GL261 GBM. **(b)** Bone marrow-derived macrophages (BMDM) were exposed to control or conditioned media (CM) from GL261 cells cultured in 1% O_2_ or normoxia. qRT-PCR results revealed upregulation of *Cd68* and *Arg1* in BMDM by CM. n=3 wells per condition, one-way ANOVA, \*\**P*<0.01, \*\*\**P*<0.001, ns, not significant. **(c)** UMAP and violin plot of TAMs show ubiquitous expression of chemokine *Ccl12*, a gene also deleted in *Ccl8/12* KO mice. **(d)** UMAP and violin plots highlight specifically expression of *Il1b* in TAMs in GL261 GBM. Note that *Il1b* is highly expressed in tumor-associated macrophages (Mac) and microglia (MG), and also in dendritic cells (DC).

**Figures S8.**
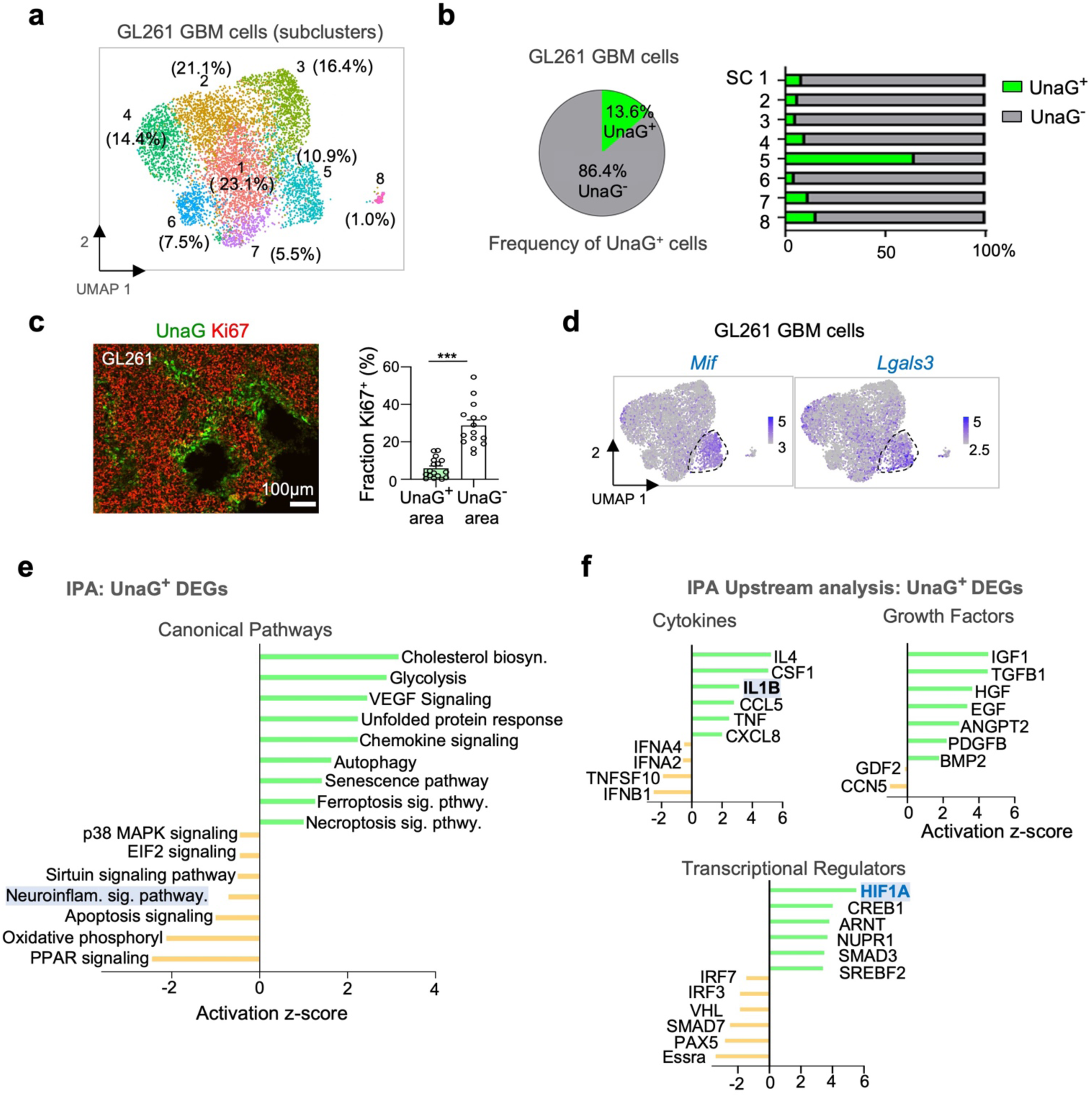
Hypoxic GBM cells display unique gene signatures showing a high degree of immune signaling. **(a)** UMAP plot of GL261 GBM cells at 4 weeks post-transplant shows 8 subclusters based on scRNA-seq data. **(b)** Among the tumor cells in GL261 GBM at 4 weeks post-transplant, 13.6% expressed high levels of UnaG mRNA. Among the 8 subclusters of tumor cells, SC-5 harbored the highest proportion of UnaG^+^ cells. **(c)** IF image of GL261 GBM for proliferation marker Ki67 and quantifications reveal that UnaG^+^ GBM cells are mainly quiescent. n=3 mice per group, unpaired t-test, \*\*\**P*<0.001. **(d)** UMAP plots show high expression of immune modulatory genes *Mif* and *Lgals3* in SC-5 (UnaG^+^) tumor cells (encircled). **(e)** Ingenuity pathway analysis (IPA) of DEGs of UnaG^+^ vs. UnaG^-^ GBM cells (defined as up or down-regulated genes with log_2_FC cut-off of 0.25). Note negative enrichment of neuroinflammation signaling pathway (highlighted in blue). **(f)** Predicted IPA upstream regulators of the DEGs in UnaG^+^ GBM cells. Note IL1B is identified as potential upstream regulator of UnaG^+^ DEGs, as is HIF1A (highlighted).

**Figure S9.**
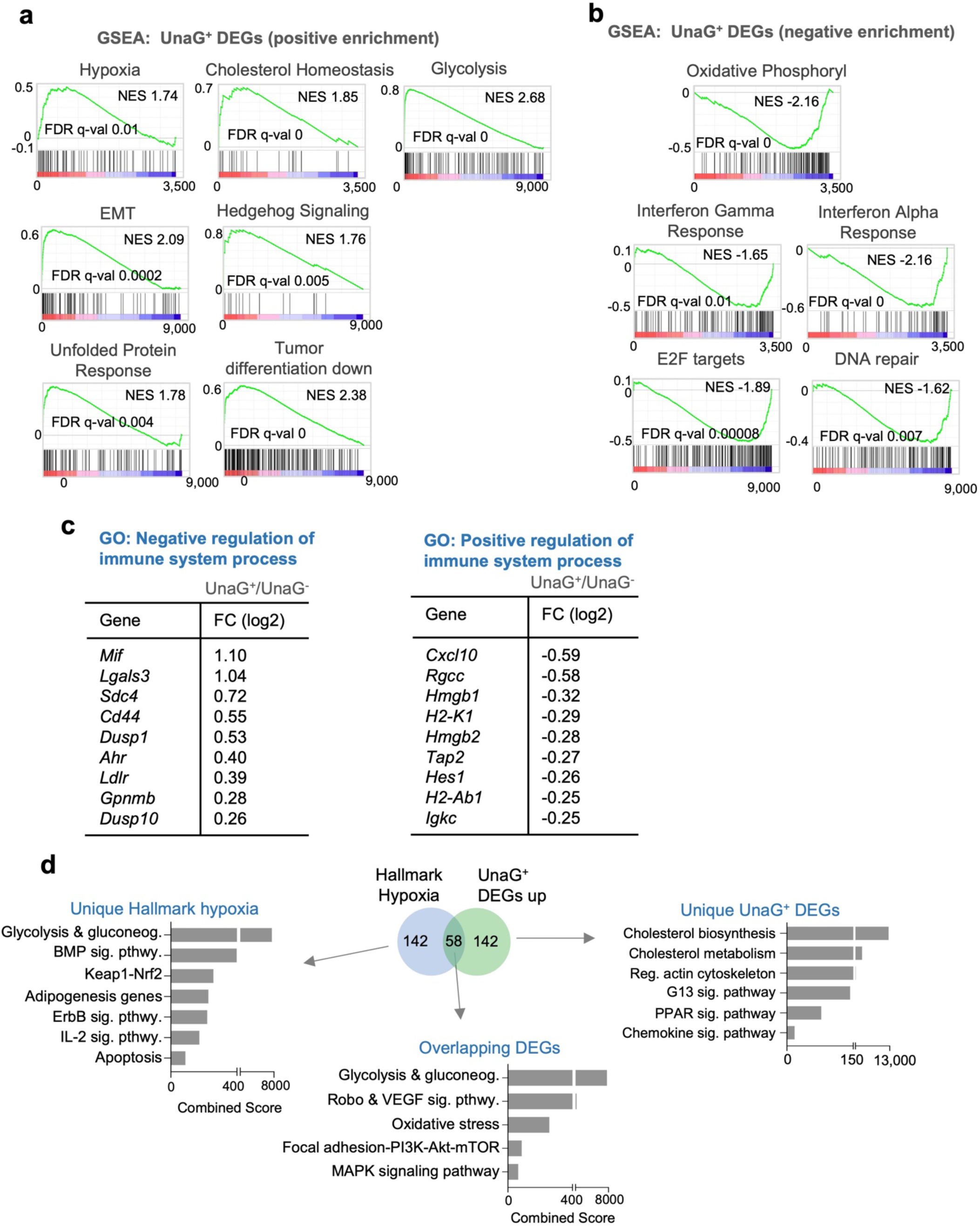
In vivo gene signatures of hypoxic GBM cells involve immune signaling pathways. **(a, b)** Gene set enrichment analysis (GSEA) of gene expression changes in UnaG^+^ GBM cells reveals top gene sets with positive enrichment (a) or negative enrichment (b) (FDR, False Discovery Rate; NES, Normalized Enrichment Score). Note negative enrichment for IFN*α* response and IFN*γ* response. **(c)** UnaG^+^ GBM cells upregulated genes associated with the GO term Negative Regulation of Immune System Process (GO:0002683), but downregulated genes of the GO term Positive regulation of Immune System Process (GO:0002684). **(d)** Venn diagram shows intersection of UnaG^+^ DEGs (200 upregulated genes) and MSigDB curated gene set ‘Hallmark Hypoxia’. Result of ENRICHR pathway enrichment analysis of the unique and shared genes are shown.

**Figure S10.**
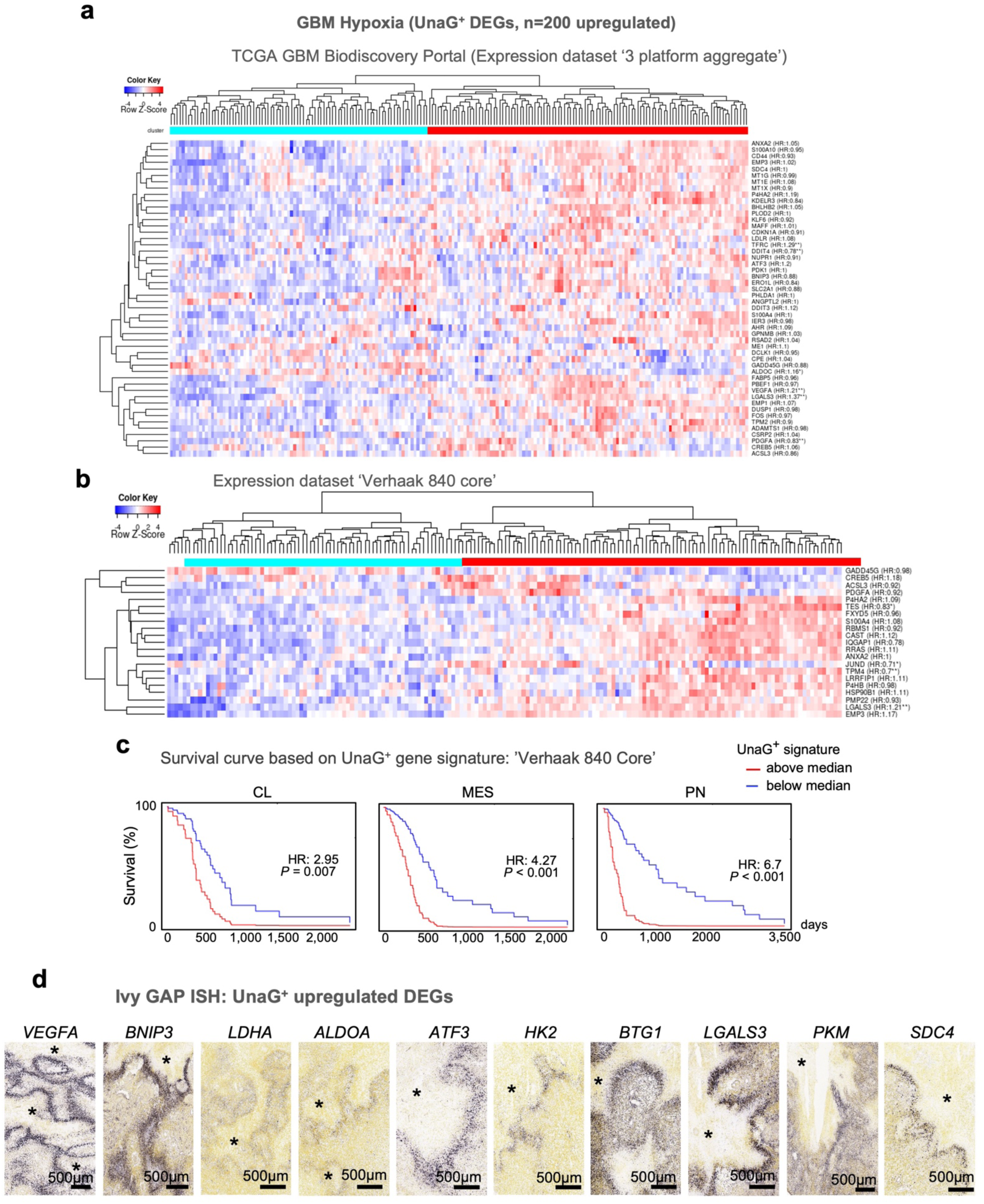
Expression of UnaG^+^ gene signature in GBM patient specimens correlates with survival. **(a, b)** Stratification of GBM patients from the GBM biodiscovery portal database according to expression scores for UnaG^+^ signature (top 200 upregulated DEGs of UnaG^+^ GBM cells). Expression datasets analyzed: 3 platform aggregates (a) and Verhaak 840 core (b). **(c)** Kaplan-Meier survival analyses of GBM patients of three different transcriptional subtypes show that high expression of UnaG^+^ gene signature is associated with shorter survival for all subtypes. Expression dataset analyzed: Verhaak 840 core. **(d)** In situ hybridization (ISH) images of Ivy GAP database show pseudopalisading expression patterns of 10 genes of the UnaG^+^ signature in patient GBM tissues. Asterisks denote hypoxic cores.

**Figure S11.**
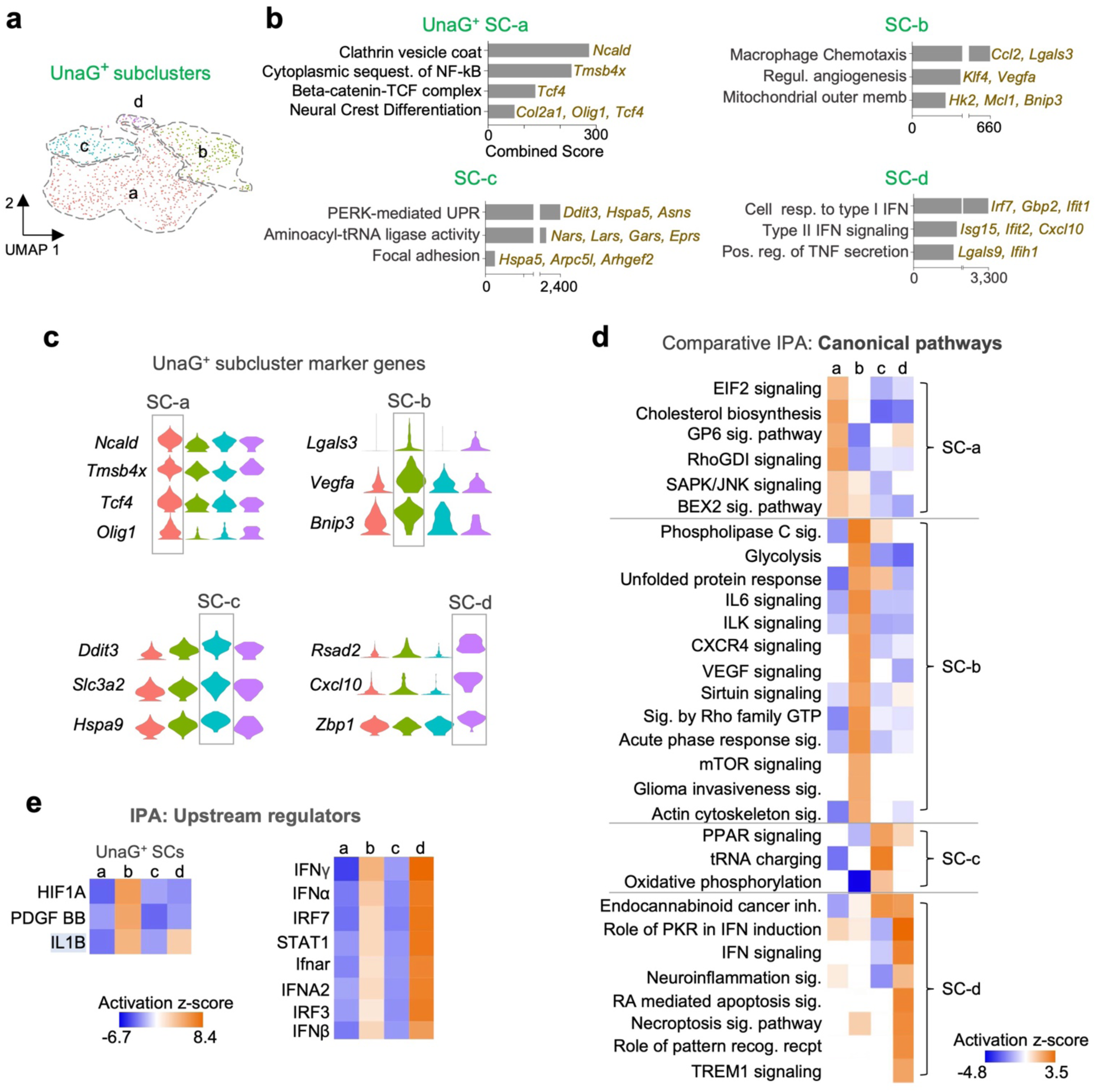
UnaG^+^ GBM cell subclusters engage distinct signaling pathways. **(a)** UMAP plot based on GL261 scRNA-seq data shows that UnaG^+^ GBM cells can be further partitioned into four subclusters (SC-a to SC-d). **(b)** ENRICHR analyses (WikiPathways 2019) show specific pathways enriched in each UnaG^+^ subcluster. **(c)** Violin plots show marker genes expression in each of the four subclusters of UnaG^+^ GBM cells. **(d)** Comparative IPA show distinct signaling pathways engaged by subclusters of UnaG^+^ GBM cells. **(e)** Comparative IPA show upstream regulators of the DEGs of UnaG^+^ subclusters.

**Figure S12.**
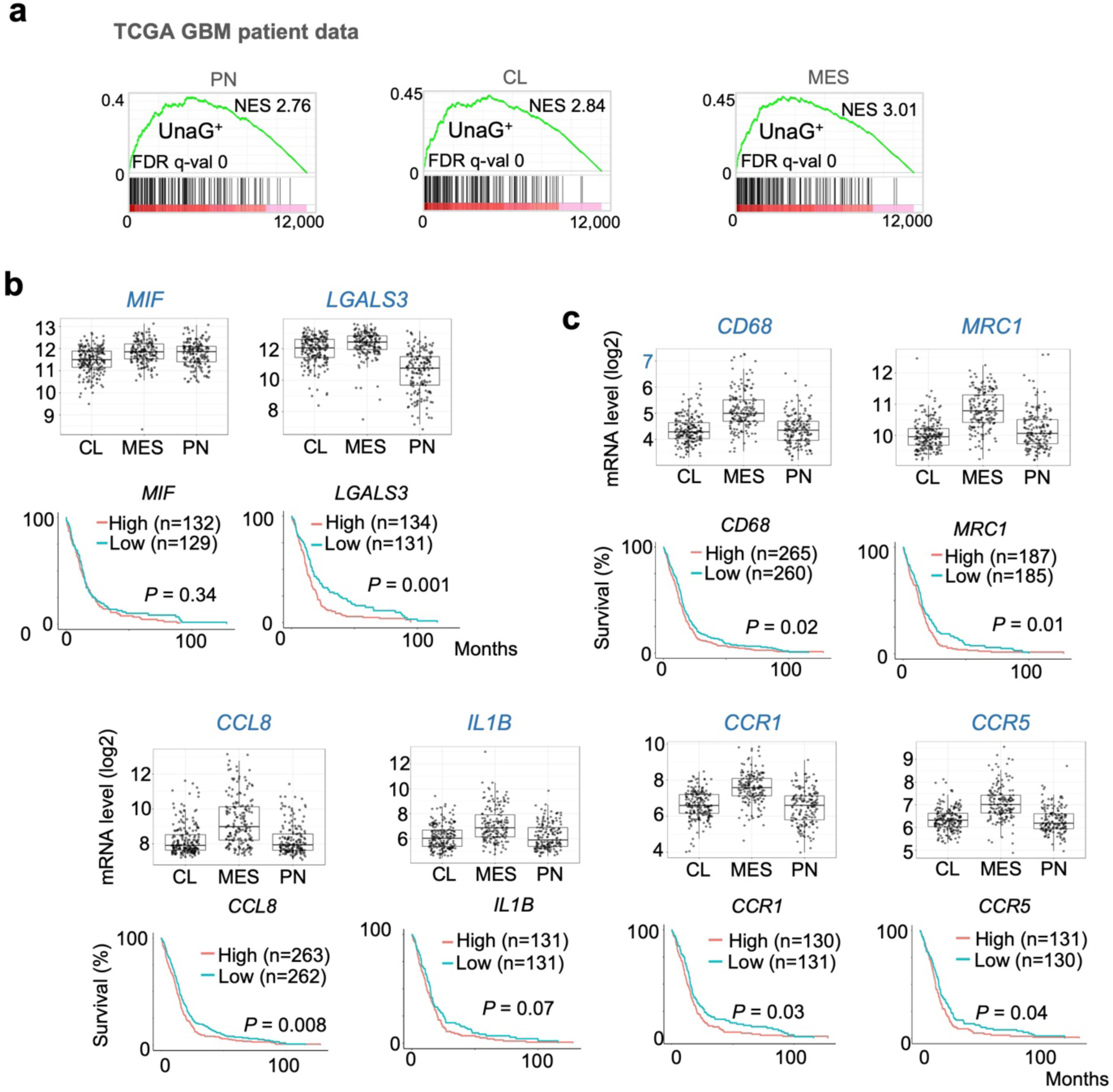
Hypoxia niche-associated genes are more represented in MES GBM subtype and linked to patient survival. **(a)** GSEA show that the UnaG^+^ gene signature (200 upregulated DEGs) is enriched in patient GBM of all three transcriptional subtypes (PN, proneural; CL, classical; MES, mesenchymal), with the MES subtype showing the highest score. For each subtype, 12,042 genes from microarray expression data were ranked. NES, normalized enrichment score; FDR, false discovery rate. **(b, c)** Analyses of TCGA GBM expression data (Gliovis, dataset: TCGA_GBM HG-U133A) show expression levels of the indicated niche genes associated with hypoxia GBM cells (b) or entrapped TAMs (c). Kaplan-Meier survival curves are constructed by dividing patients by median expression into high and low expressor cohorts. Overall, all these niche genes show higher expression in the MES subtype, and they predict worse survival (except for *MIF*).

